# ApoE4 disrupts intracellular trafficking and iron homeostasis in an improved iPSC-based model of human brain endothelial cells

**DOI:** 10.1101/2024.09.03.610802

**Authors:** Luisa Bell, Nadine Stokar-Regenscheit, Claire Simonneau, Angélique Augustin, Barbara Höllbacher, Lia D’Abate, Joanna Ficek-Pascual, Kim Schneider, Desiree Von Tell, Chiara Zanini, Christelle Zundel, Sabrina Golling, Christine Becker, Alex Odermatt, Lynette Foo, Martina Pigoni, Roberto Villaseñor

## Abstract

Transferrin receptor in brain endothelial cells can deliver therapeutic antibodies to the brain via transcytosis across the blood-brain barrier. Whether receptor transport remains intact in Alzheimer’s disease is still a major open question. Here, we investigated whether apolipoprotein E4 (ApoE4), the major genetic risk factor for Alzheimer’s disease, altered intracellular transport in human brain endothelial cells. To achieve this, we first developed an optimized protocol for induced pluripotent stem cells based on a defined <u>c</u>hemical cocktail and <u>e</u>xtracellular-matrix support to differentiate <u>b</u>rain <u>e</u>ndothelial <u>c</u>ells (iCE-BECs). Multi-omic profiling and functional transport assays showed that iCE-BECs have a brain endothelial gene signature and recapitulate receptor-mediated transcytosis of a clinically validated Brainshuttle^TM^ antibody against transferrin receptor. Engineered iCE-BECs homozygous for ApoE4 had altered spatiotemporal organization of early endosomes, increased transferrin receptor expression and reduced cytoplasmic iron. Our data revealed that ApoE4 can impact intracellular transport and iron homeostasis at the BBB in a cell-autonomous manner. This finding could be relevant for the brain delivery of therapeutic antibodies for Alzheimer’s disease.

## Introduction

The blood-brain barrier (BBB) is formed by brain endothelial cells, pericytes and astrocytes organized into a neurovascular unit which regulates the exchange of proteins between blood circulation and brain parenchyma via receptor-mediated transcytosis ^1–7^. The transferrin receptor (TfR1) is one of the best characterized receptors involved in transcytosis and is validated as a target to deliver therapeutic antibodies to the brain parenchyma ^8–10^. Intracellular sorting in endosomes determines whether a receptor undergoes transcytosis or is instead transported to lysosomes for degradation ^11–13^. Whether intracellular trafficking in brain endothelial cells is altered in disease conditions and to what extent this impacts transcytosis across the BBB remain outstanding open questions.

There is a wealth of evidence that documents the impact of disease conditions on BBB paracellular permeability ^14,15^. For example, it is well established that ApoE4, the major genetic risk factor for sporadic Alzheimer’s disease (AD), increases paracellular permeability of the BBB ^16–21^. On the other hand, disease-specific changes to intracellular transport and/or transcytosis across the BBB are still relatively unexplored. A recent transcriptomic analysis of the vasculature of AD brain tissue showed substantial changes in gene expression levels associated with intracellular trafficking, including down-regulation of TfR in brain capillaries ^22^. However, disease subgroups or risk factor genotypes could further affect TfR expression and trafficking. For instance, it is known that ApoE4 expression in mice led to upregulation of TfR expression in brain endothelial cells ^23^. This highlights the need for a systematic approach to evaluate how specific risk factors affect transport across the BBB.

Human stem-cell based models using brain endothelial cells are a powerful tool to investigate how disease-related conditions might affect BBB integrity ^24,25^. However, earlier studies used models with an overt epithelial signature that lacked expression of genes required for endothelial function ^26^. Since the mechanisms of intracellular transport in endothelial and epithelial cells are different ^13,27–29^, inducible pluripotent stem cells (iPSC)-based models with epithelial identity are not suitable to investigate the regulation of transcytosis across the BBB. To address this, we developed a protocol to differentiate endothelial cells with brain-specific identity from iPSC. We showed that these cells expressed key markers of brain endothelial cells and recapitulated receptor-mediated transcytosis of a clinically validated TfR-dependent Brainshuttle^TM^ antibody. Using this differentiation protocol on genetically engineered iPSC lines, we found that ApoE4 regulated the organization and dynamics of the early endosome network in a cell-autonomous manner in brain endothelial cells. Finally, we showed that ApoE4 also dysregulated iron homeostasis, triggering overexpression of TfR in brain endothelial cells. Our data highlights the utility of relevant human *in vitro* models of brain endothelial cells to investigate how disease risk factors affect intracellular transport and reveal a new role for ApoE4 in the regulation of iron metabolism at the BBB.

## Results

### Characterization and validation of a novel protocol for differentiation of brain endothelial cells from iPSC to investigate protein transport

To generate human brain endothelial cells from iPSC, we first triggered differentiation into endothelial cells via mesoderm induction as previously described ^30^. After endothelial differentiation, we treated these cells with a defined chemical cocktail that modulates key pathways associated with induction of BBB properties during development: Wnt, cAMP and TGF-β b (Figure 1A & methods for details) ^31^. In addition to this chemical cocktail, we replated the cells in vitronectin, an extracellular matrix (ECM) component with a key role in the regulation of transcytosis in mice *in vivo* ^32^. We termed the cells generated by this protocol iCE-BECs (Inducible differentiation via <u>C</u>hemical cocktail and <u>E</u>xtracellular matrix support for <u>B</u>rain <u>E</u>ndothelial <u>C</u>ells). Previous work showed that inhibition of TGF-β signaling by RepSox enhanced barrier properties in inducible Endothelial cells (iECs) ^33^. We therefore asked whether iCE-BECs improved barrier properties beyond those triggered by RepSox alone. To this end, we compared iCE-BECs with iECs (Figure 1A) and with iECs treated with RepSox (iEC-rep, Figure 1A). Cell lines derived from all three protocols expressed endothelial markers PECAM1, VE-Cadherin, and Claudin-5 (Figure 1B). However, both Claudin-5 expression and the number of cells expressing Claudin-5 was higher in iCE-BECs compared to iECs and iEC-rep after 14 days of differentiation (Figure 1B, Figure S1A and S1B). Additionally, the iCE-BEC protocol resulted in a higher yield of PECAM1-positive cells after 11 days of differentiation (10% in iECs vs 28% in iCE-BECs PECAM1-positive cells, Figure S1C and S1D). The higher number of cells expressing Claudin-5 and PECAM1 shows that the iCE-BEC protocol generally improved efficiency of differentiation. To better characterize the cell identity and heterogeneity of iCE-BECs, we performed single-cell RNA sequencing on iECs, iEC-rep and iCE-BECs. Analysis was performed at day 14 in culture and data were integrated to assess the composition of the cell population obtained with each protocol (Figure 1C). Whole transcriptome analysis showed that endothelial markers (e.g. *PECAM1, CDH5, CLDN5*) were enriched in the cluster associated with the iCE-BEC protocol, whereas mural cell markers (e.g. *MYL9, TAGLN, ACTA2*) were enriched in iECs and iEC-rep protocols (Figure 1D, Figure S1E). To characterize the cell identity of iCE-BECs, we evaluated the expression of a recently described list of genes ^26^ (see Table S1) associated with an endothelial transcriptomic signature (Positive PC1 loading) or an epithelial transcriptomic identity (Negative PC1 loading). Endothelial signature genes were enriched in iCE-BECs, whereas epithelial signature genes were mildly expressed across the three protocols (Figure 1E). Overall, the transcriptomic data showed that iCE-BECs have endothelial identity and have a more homogeneous cell population compared to iECs.

**Figure 1.**
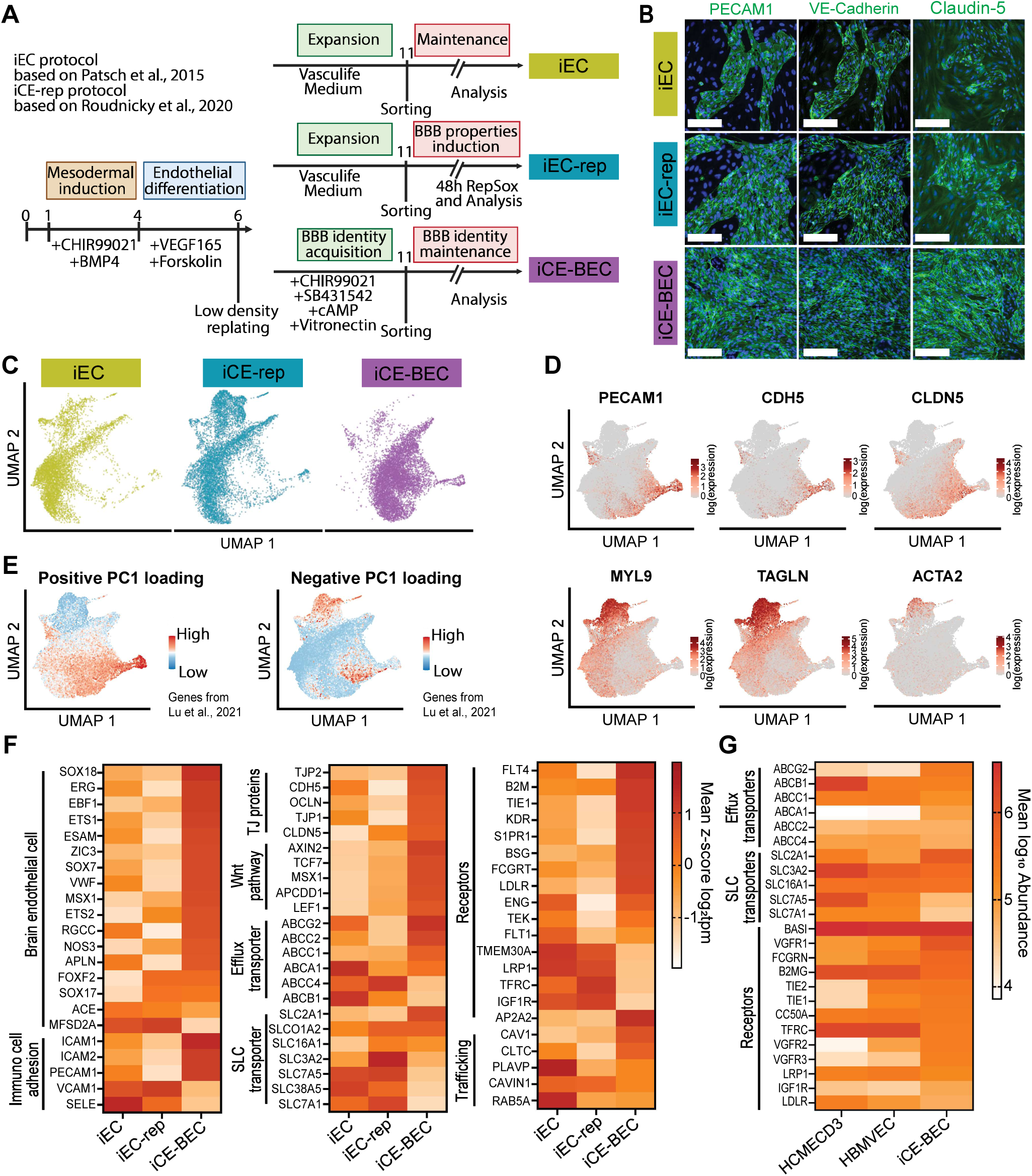
Differentiation of Brain Endothelial Cells from Induced pluripotent stem cells by a Chemical cocktail and ECM support (iCE-BECs). **A,** Schematic of the protocols used for iPSC differentiation. All protocols start from mesodermal induction of iPSC lines. Induced endothelial cells (iECs) are differentiated as described in ^30^; Induced endothelial cells with RepSox (iEC-rep) is used to enhance barrier properties of endothelial cells as described in ^33^; Inducible pluripotent stem cells via <u>C</u>hemical cocktail and <u>E</u>xtracellular matrix support-<u>B</u>rain <u>E</u>ndothelial <u>C</u>ells (iCE-BECs) are generated by a new protocol based on the simultaneous inhibition of TGF-β (SB431542), activation of cAMP and Wnt signaling (CHIR99021), and growth in vitronectin. **B,** Representative fluorescence images after immunostaining with endothelial-specific markers of cells generated using the protocols summarized in Fig. 1A. Cells are pseudo-colored showing PECAM1, VE-Cadherin or Claudin-5 in green, and DAPI-stained nuclei in blue. Scale bar, 200 µm. **C,** Integrated UMAP plot of scRNA-seq analysis, performed on the endothelial cells generated using the protocols summarized in Fig. 1A. Data are generated from 3 independent differentiations per condition. **D,** Feature plots showing normalized expression of marker genes of endothelial (*PECAM1, CDH5* and *Claudin-5*) and mural (*MYL9, TAGLN* and *ACTA2*) markers, plotted on the UMAP from Fig. 1C. **E,** Feature plots showing expression of modules “positive PC1 loading” and “negative PC1 loading” using the gene list described in Lu, Houghton, Magdeldin, Durán, Minotti, Snead, Sproul, Nguyen, Xiang, Fine, Rosenwaks, Studer, Rafii, Agalliu, Redmond and Lis ^26^ (see Table S1), plotted on the UMAP from 1C. **F,** Bulk RNA-Seq heatmaps showing expression of genes for main brain endothelial, immune cell adhesion, tight junction (TJ) proteins, Wnt pathway, efflux and Solute Carrier (SLC) transporters, receptors and trafficking markers across the three different differentiation protocols for iECs, iEC-rep, and iCE-BECs. Values are expressed as mean z-score log_2_tpm, with three independent differentiations per condition. **G,** Whole cell proteomics heatmap showing expression of main endothelial and brain endothelial receptors and transporters comparing immortalized human brain endothelial cells (HCMEC/D3, n=3), primary human microvascular brain endothelial cells (HBMVEC, n=6) and iCE-BECs (n=4 independent differentiations). Values are expressed as mean log_10_ Abundance.

Next, we performed bulk RNA sequencing analysis to evaluate whether iCE-BECs have an improved brain-specific signature. Endothelial and brain endothelial cell markers were upregulated in iCE-BECs compared to iECs and iEC-rep (Figure 1F), while epithelial and mural cell markers were not detected or strongly downregulated (Figure S2). iCE-BECs showed an upregulation of *MSX1, ZIC3, EBF1* and *APLN*, which are genes specifically enriched in brain endothelial cells in mice ^34–36^. Additionally, iCE-BECs had an upregulation of genes associated with BBB function, including tight junction genes (*CLDN5, TJP1, TJP2, OCLN, CDH5*), solute carriers (*SLC2A1, SLCO1A2, SLC16A1*), efflux transporters (*ABCG2, ABCC2*) (Figure 1F), genes involved in immune cell adhesion (*ICAM1, ICAM2, PECAM1)* and downstream effectors of Wnt signaling (*LEF1, TCF7, AXIN2, APCDD1*), ^37^. These data confirm that iCE-BECs exhibit a brain identity signature and express key genes essential for brain endothelial function.

To investigate the applicability of iCE-BECs for transport assays, we measured the expression of transporters and receptors with whole cell proteomics. We compared this data to the proteome of both primary human brain microvascular endothelial cells (HBMVECs) and the widely used immortalized brain endothelial cell line HCMECD3. We observed small changes in specific receptors that were upregulated (SLC2A1, ABCA1, ABCG2) or downregulated (SLC7A1, SLC7A5) in iCE-BECs. However, the proteomics analysis showed an overall similar expression of transporters and receptors between the three cell lines (Figure 1G). Taken together, transcriptomics and proteomics data demonstrated that iCE-BECs are a homogeneous population of cells with human brain endothelial identity and express proteins relevant for the *in vitro* assessment of transcytosis across the BBB.

### iCE-BECs restrict transport of large molecules and recapitulate receptor-mediated transcytosis

Next, we assessed whether the iCE-BEC protocol improved paracellular permeability compared to cells treated with RepSox. Since the main context of use for this model is to investigate mechanisms of antibody transport, we focused our permeability measurements on dextrans of different molecular weights (3, 40, and 70 kDa) on transwell chambers. We found that iCE-BECs had a 75% reduction in permeability for dextran across different sizes compared to both iECs and iEC-rep (Figure 2A). Since iCE-BECs grew in a continuous monolayer in the transwell filter (Figure 2B) their observed decreased permeability is not due to the formation of multiple pseudo-stratified cell layers in the filter.

**Figure 2.**
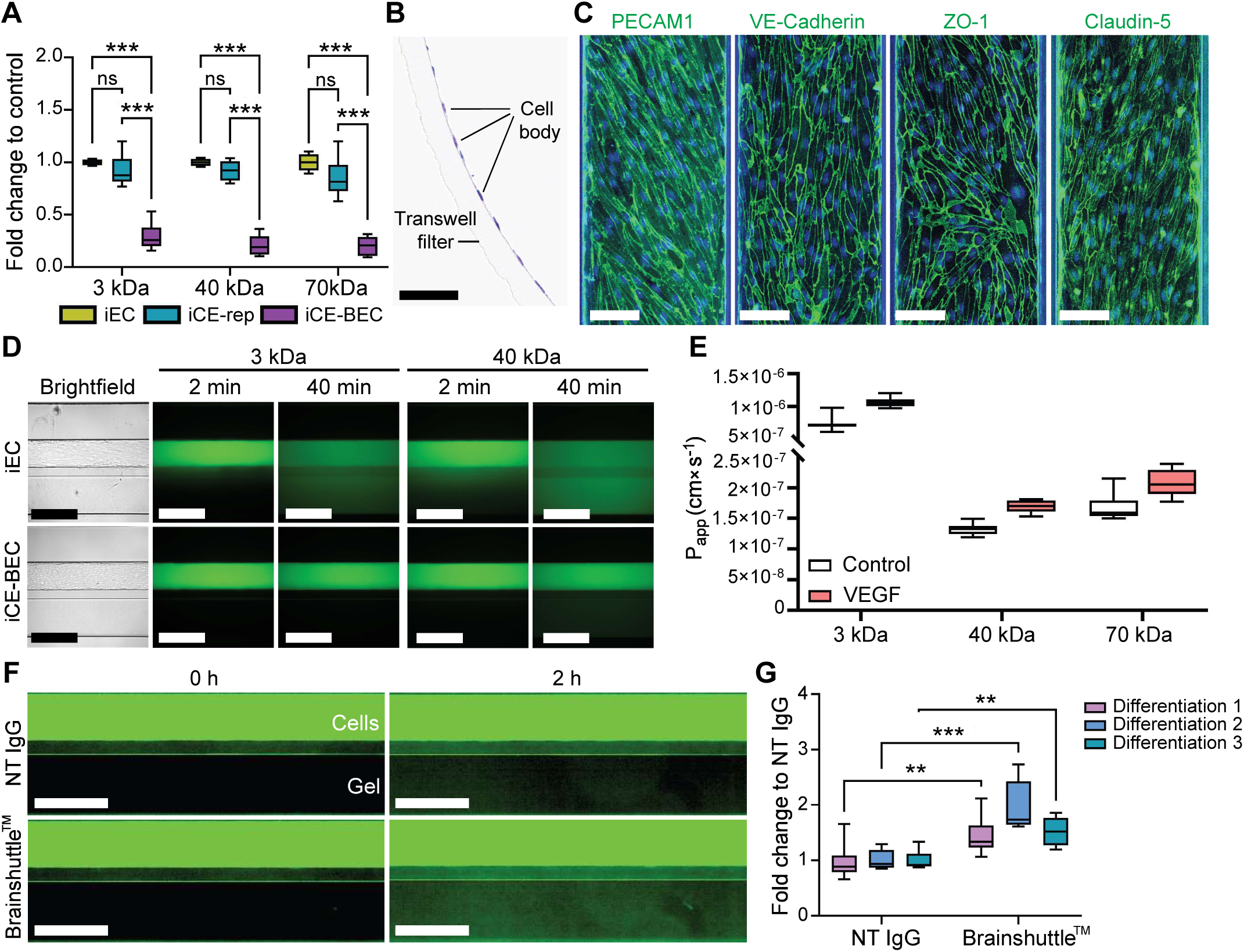
iCE-BECs have lower permeability to large molecules compared to iECs and recapitulate receptor-mediated transcytosis of Brainshuttle™ antibodies. **A,** Quantification of relative apparent permeability (P_app_) for dextrans of different molecular weights of cells generated with the protocols described in Fig. 1A using a transwell system. Values were normalized to the apparent permeability of iECs. Graph shows boxplots with interquartile ranges and median. Lines show the 5th and 95th percentiles, data from n = 4 independent differentiations with 8 technical replicates per condition. Differences in apparent permeability are statistically significant as evaluated by Two-way ANOVA with Sidak multiple comparisons. ns, not statistically significant; ***, p <0.001. **B,** Haematoxylin and Eosin staining of iCE-BECs grown on a transwell filter showing a cell monolayer. Scale bar, 50 µm. **C,** Representative fluorescence images after immunostaining with endothelial-specific markers pseudo-colored showing PECAM1, VE-Cadherin, ZO-1 and Claudin-5 of iCE-BECs grown in a MIMETAS OrganoPlate® 2-lane 96, Scale bar, 100 µm. **D,** Representative bright field (grey left panels) and fluorescence images of MIMETAS OrganoPlate® 2-lane chambers comparing iEC (top) and iCE-BECs (bottom) after incubation with 3 kDa or 40 kDa dextran for 2 or 40 minutes. The ratio between fluorescent intensity between the upper (vascular cell compartment) and lower gel chamber is used for the estimation of apparent permeability, Scale bar, 500 µm. **E,** Quantification of apparent permeability (P_app_) of iCE-BECs to 3, 40 or 70 kDa dextran in basal conditions or after stimulation with 200 ng/mL VEGF-A for 24 hours. Graph shows boxplots with interquartile ranges and median. Lines show the 5th and 95th percentiles, data from one differentiation with at least 7 technical replicates per condition. **F,** Representative fluorescent images of iCE-BECs grown on MIMETAS OrganoPlate® 2-lane chambers comparing iEC (top) and iCE-BECs (bottom) after incubation with 200 nM non-targeting IgG (NT IgG) or a Brainshuttle™ antibody. Images were acquired immediately after incubation (0 hours) or after 2 hours. Scale bar, 500 µm. **G,** Quantification of relative antibody transcytosis across iCE-BECs (see Methods for details). Values are normalized relative to the NT-IgG condition. Graph shows boxplots with interquartile ranges and median. Lines show the 5th and 95th percentiles, data from 3 independent differentiations, each at least 7 technical replicates per condition. Differences in transcytosis are statistically significant as evaluated by Two-way ANOVA with Sidak multiple comparisons. **, p <0.01; ***, p <0.001.

To further characterize iCE-BEC permeability we used a microfluidic platform (MIMETAS OrganoPlate® 2-lane 96) previously established to investigate BBB transport ^38,39^. As expected, iCE-BECs grown on the MIMETAS platform expressed endothelial markers and tight junction proteins (Figure 2C). Permeability to fluorescently-labeled dextrans was measured over time (Figure 2D) and was substantially reduced compared to iECs and a chip without any cells (Figure S3A). We estimated that iCE-BECs had an apparent permeability in the order of 10^-7^ cm/s^2^ for 40 kDa dextran (Figure 2E, see methods for details). Treatment of iCE-BECs with VEGF increased the permeability to dextran, further confirming the endothelial identity of the cells (Figure 2E). Importantly, we reproduced the same apparent permeability range when applying the iCE-BEC differentiation protocol to 4 different iPSC lines (Figure S3B). This result shows that the iCE-BEC protocol is not restricted to a single parental line but could be broadly applicable to multiple iPSC lines.

Since iCE-BECs restrict the permeability of large molecules in the MIMETAS platform, we evaluated receptor-mediated transcytosis in this model. To do this, we used the Brainshuttle^TM^, a validated TfR1 binder fused to an antibody cargo that effectively crosses the BBB ^10,40^. We found that the Brainshuttle^TM^ construct showed an almost 2-fold higher transport rate compared to a non-targeting IgG (Figure 2F, G; Figure S3C, D). This result is in agreement with previously reported values for the same antibody in BBB organoids ^41^. Importantly, we confirmed that the extent of Brainshuttle^TM^ transcytosis was reproducible across 3 different iPSC lines differentiated with the iCE-BEC protocol (Figure S3E). These data show that iCE-BECs recapitulate key features of the BBB including low permeability to large molecules and receptor-mediated transcytosis. Together, our data show that iCE-BECs 1) have endothelial identity, 2) show a brain-endothelial signature and 3) recapitulate receptor-mediated transcytosis of antibodies. Therefore, we concluded that iCE-BECs are a suitable human *in vitro* system to investigate the regulation of intracellular transport across brain endothelial cells and, more specifically, the impact of disease risk factors on transport.

### ApoE4 alters endosome morphology and dynamics in iCE-BECs

ApoE4 is the major risk factor for sporadic Alzheimer’s disease ^20,42,43^. Multiple studies show that ApoE4 can affect BBB function, likely via non-cell autonomous mechanisms ^19,23,44^.

Nevertheless, it is unclear whether ApoE4 affects brain endothelial cells in a cell-autonomous manner ^45^. To address this, we used iPSC lines with either a homozygous ApoE3 (Bioni037-A) or homozygous ApoE4 (Bioni037-A4) gene variant and differentiated them into iCE-BECs. The homozygous ApoE3 line was generated from fibroblasts of a healthy individual genetic variant ^46^ and the isogenic homozygous ApoE4 was derived by an insertion of the point mutations required for the ApoE4 gene variant (see methods for details). Importantly, ApoE4 did not change the proliferation rate of iPSCs (Figure S4A) and did not impair the differentiation process as evidenced by expression of endothelial and tight junction genes (Figure 3A, Figure S4B). The mRNA and protein expression levels of *APOE* were lower in ApoE4 compared to ApoE3 iCE-BECs (Figure 3B, C). Such lower expression of ApoE is in agreement with previous data obtained from ApoE4 astrocytes ^47,48^. These data suggest that ApoE4 iCE-BECs can be used to investigate cell autonomous effects of ApoE4.

**Figure 3.**
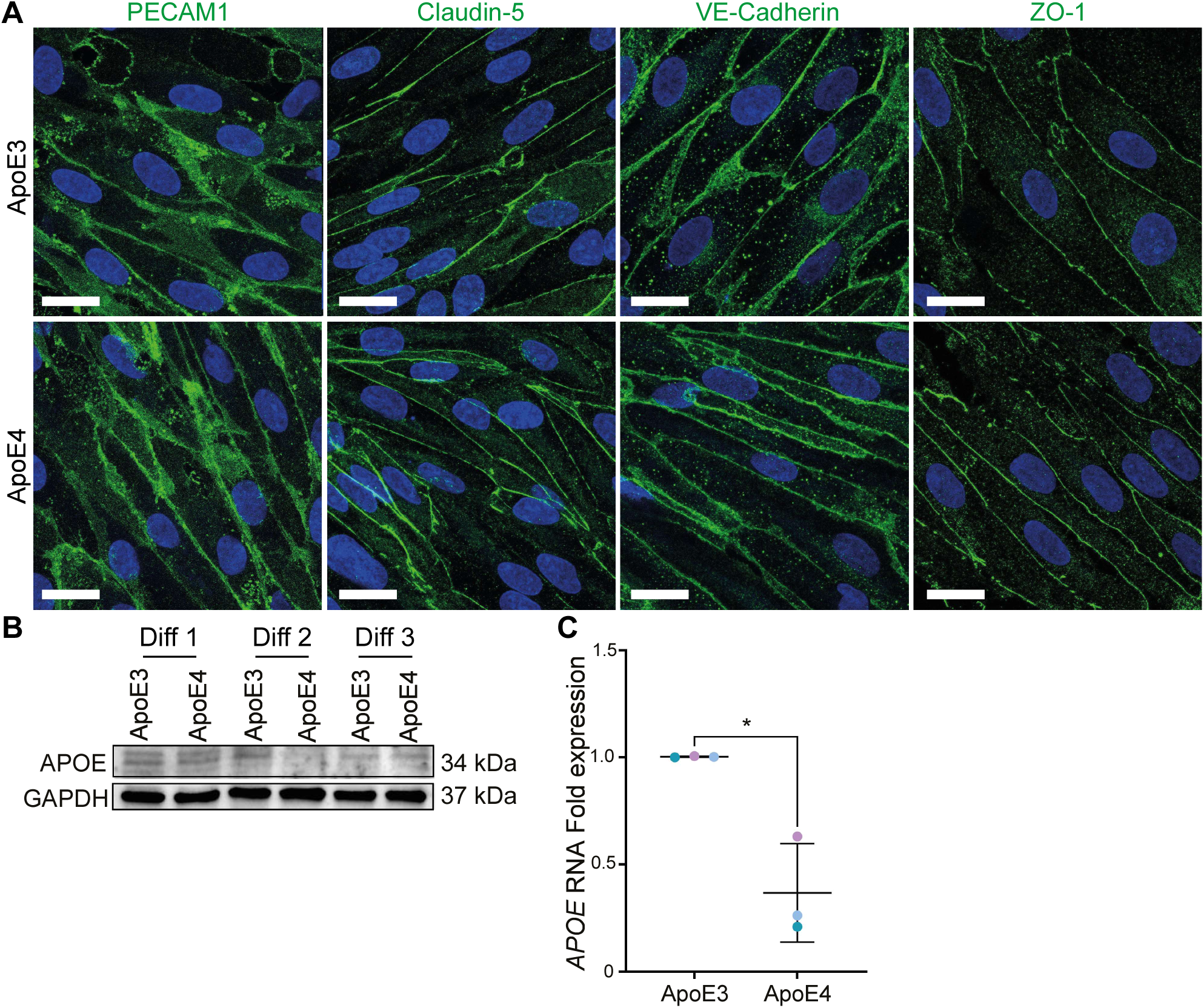
ApoE4 genetic variant does not alter differentiation of iCE-BECs. **A,** Representative fluorescence images after immunostaining with endothelial-specific markers in iCE-BECs with ApoE3 or ApoE4 genetic variant. Cells are pseudo-colored showing PECAM1, VE-Cadherin, ZO-1 or Claudin-5 in green, and DAPI-stained nuclei in blue. Scale bar, 20 µm. **B,** Representative Western Blot image showing ApoE protein expression in iCE-BECs with ApoE3 or ApoE4 genetic variant. Data come from three independent differentiations (Diff). **C,** Quantification of relative ApoE mRNA expression by quantitative PCR. Graph shows mean ± SD of n = 3 independent differentiations with 3 technical replicates per experiment. Points represent independent differentiations. Differences in ApoE mRNA expression are statistically different between ApoE genetic variant as evaluated by the Student’s t test. *, p <0.05.

Recent work showed that ApoE4 alters multiple endocytosis mechanisms *in vivo* ^23,49^. Therefore, we analyzed the organization of endosomes in ApoE4 iCE-BECs. We found that the number of early endosomes (labeled by early endosomal antigen 1, EEA1) was higher in ApoE4 compared to ApoE3 iCE-BECs (Figure 4A, C). In addition, the total amount of EEA1 per endosome (estimated by the mean integral vesicular intensity) was also higher in ApoE4 compared to ApoE3 iCE-BECs (Figure 4B). To confirm the changes to the endosomal network with an orthogonal method, we analyzed iCE-BECs with transmission electron microscopy (TEM) to evaluate the ultrastructural morphology of endosomes. We observed that ApoE4 iCE-BECs had an increase in endosome number and size compared to ApoE3 cells (Figure 4A-D). We hypothesized that such changes to endosome number, size and EEA1 amount in ApoE4 iCE-BECs arise from changes to the endosome fusion/fission equilibrium ^50^. To test this hypothesis, we used live-cell imaging to visualize the dynamics of fluorescent transferrin, a well-established marker for early and recycling endosomes. In agreement with previous data on mouse brain endothelial cells ^29^, we observed frequent events of sorting tubule biogenesis and vesicle fission in ApoE3 iCE-BECs (Figure 4E, Video S1, 2). In contrast, the number of sorting tubules was substantially reduced in ApoE4 iCE-BECs (Figure 4F). Together, these data show that ApoE4 impacts sorting tubule formation and fission, which are expected to increase both endosome number and size.

**Figure 4.**
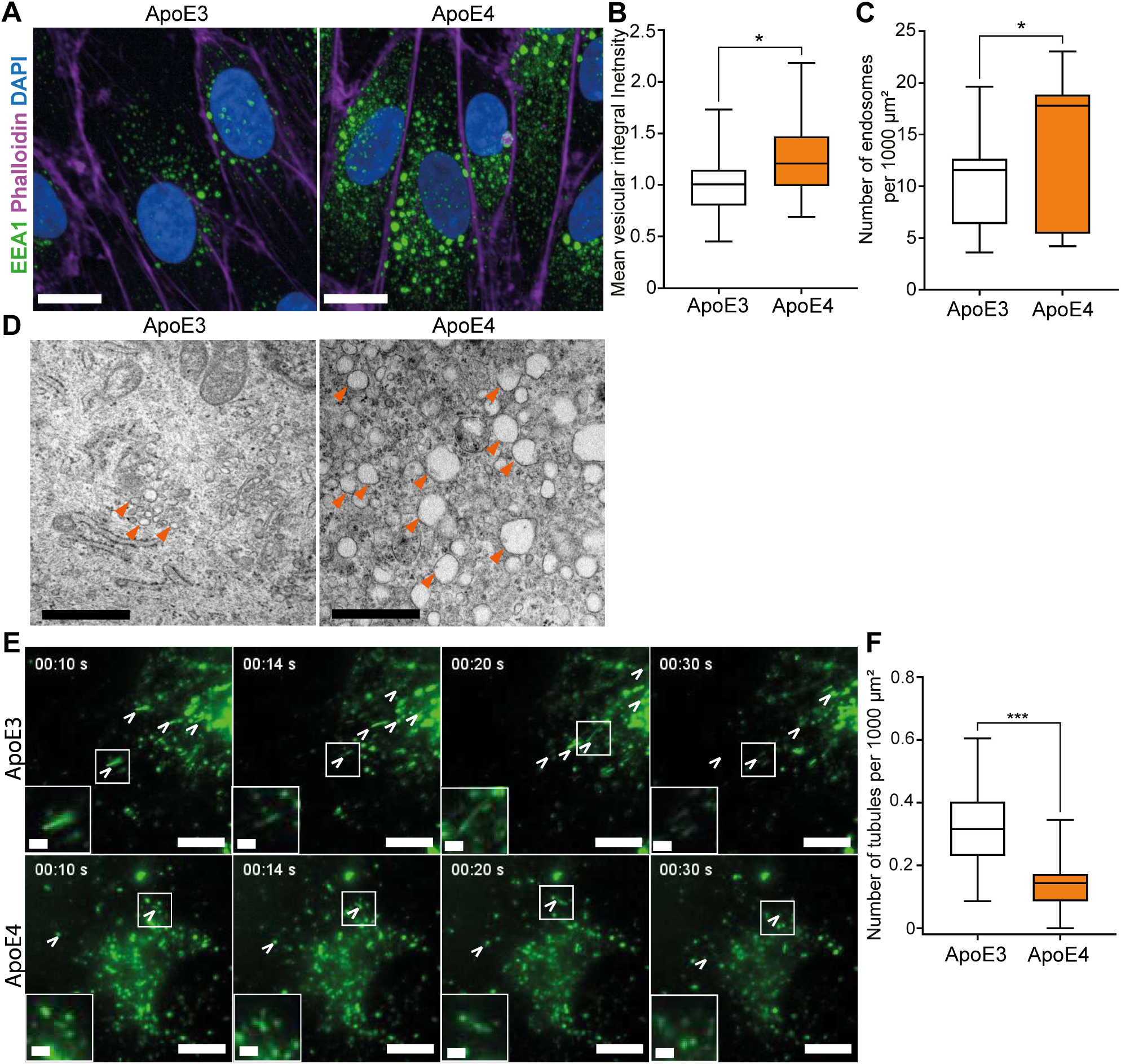
ApoE4 alters early endosomes and reduces sorting tubule biogenesis in iCE-BECs. **A,** Representative maximum intensity projections of fluorescence confocal images of iCE-BECs with ApoE3 or ApoE4 genetic variant after immunostaining with the marker of early endosomes (EEA1). Cells are pseudo-colored showing EEA1 in green, Phalloidin in magenta and DAPI-stained nuclei in blue. Scale bar, 10 µm. **B-C,** Quantification of mean integral vesicular intensity of EEA1 normalized to Phalloidin-postive area **(B)** or number of EEA1-positive endosomes per 1000 μm^2^ Phalloidin-positive area **(C).** Values in B were normalized to ApoE3 conditions for each experiment. Graph shows boxplots with interquartile ranges and median. Lines show the 5th and 95th percentiles. Per experiment, 10 images were acquired with approximately 200 cells per condition, data come n = 3 independent differentiations. Differences in EEA1 intensity and number of EEA1-positive endosomes are statistically significant as evaluated by the Mann-Whitney-U test. *, p <0.05. **D,** Representative transmission electron microscopy images of ApoE3 or ApoE4 iCE-BECs. Arrowheads point to endosomes. Scale bar, 1 µm. **E,** Representative images of transferrin (green) intracellular transport in live iCE-BECs with ApoE gene variants after incubation with 25 µg/ml of Transferrin-Alexafluor488 for 3 hours. The time stamp shows the elapsed time in seconds after the acquisition of the first image. Arrowheads point to individual sorting tubules across time frames. Inserts depict a zoom in on selected sorting tubules. Scale bar, 10 µm, inserts 2 µm. **F,** Quantification of the mean number of sorting tubules per 1000 µm^2^ occurring in one minute. Graph shows boxplots with interquartile ranges and median. Lines show the 5th and 95th percentiles, data come from n = 3 independent differentiations with 20 videos per experiment and per genetic variant. Differences in the number of tubules are statistically significant as evaluated by the Student’s t test. ***, p < 0.001.

### ApoE4 increases transferrin receptor expression without altering its transport rate in iCE-BECs

We next asked whether the changes in the organization of the early endosome network led to functional consequences in intracellular transport rates. To evaluate this, we followed the uptake and recycling of fluorescently labeled transferrin in iCE-BECs using a continuous pulse or a pulse-chase experimental design, as previously described (Figure S5A, B) ^29^. ApoE4 iCE-BECs had a 50% higher total amount of internalized transferrin compared to ApoE3 iCE-BECs (Figure 5A). This increase in total internalized transferrin was confirmed by flow cytometry (Figure S5C). However, the rate of transferrin internalization was similar between ApoE4 and ApoE3 iCE-BECs (Figure 5C). Similarly, the total amount of transferrin after a 20 minute pulse was higher in ApoE4 compared to ApoE3 iCE-BECs (Figure 5B), but the recycling rate was similar between the two genetic variants (Figure 5D). An increased transferrin uptake capacity without changes to the internalization and recycling rates could be explained by the upregulation of TfR expression. Indeed, we found that both mRNA and protein TfR expression were upregulated in ApoE4 compared to ApoE3 iCE-BECs (Figure 5E-H). These data show that ApoE4 leads to TfR upregulation but does not alter intracellular transport rates in iCE-BECs. To confirm that these results are not caused by the specific genetic background of the parental cell line, we assessed the effect of the ApoE4 gene variant on a different iPSC line. In this case, the parental ApoE4 line was generated by reprogramming of human bone marrow CD34-positive mononuclear cells carrying ApoE4 in both alleles (Alstem iPS16); the isogenic homozygous ApoE3 (Alstem iPS26) was derived by genome editing of the ApoE allele (see methods for details). Importantly, the same phenotypes (i.e. enlarged early endosome network, increased transferrin uptake and TfR expression) were consistently reproduced in this genetic background (Figure S6).

**Figure 5.**
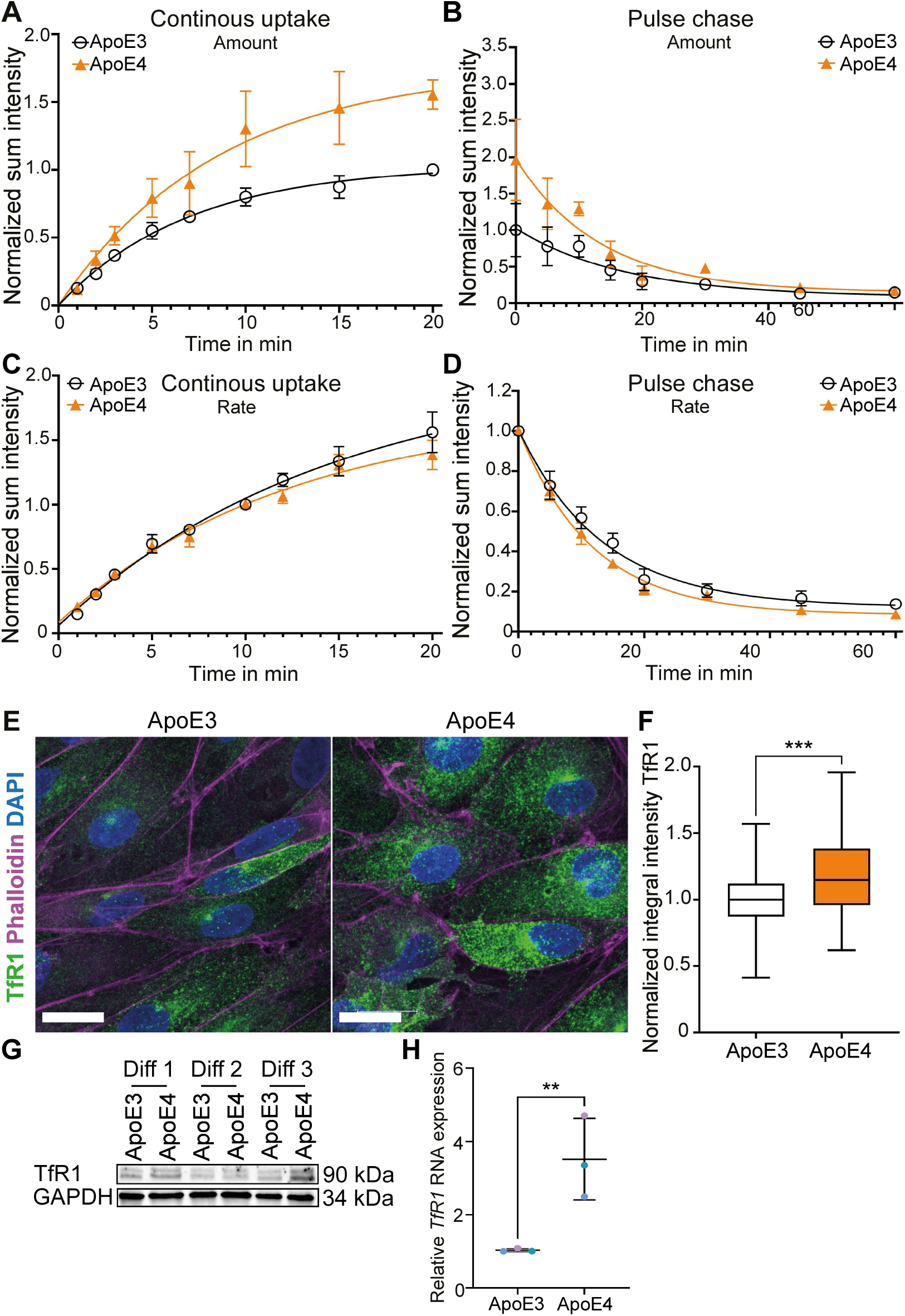
ApoE4 iCE-BECs have increased TfR1 expression but no changes to trafficking kinetics. **A-D,** Time courses of continuous uptake **(A)** or pulse-chase **(B)** assays showing total vesicular intensity of transferrin in ApoE3 or ApoE4 iCE-BECs. Intensity values were divided by the mean of the transferrin intensity at 20 min **(A)** or immediately after the pulse **(B)** in ApoE3 iCE-BECs. Points show the average and error bars show the SEM from 50 images per experiment in 4 **(A)** or 6 **(B)** independent differentiations. Lines show the best exponential fit for the experimental data. For illustration of the transferrin kinetics, graphs in **C** and **D** were normalized to the transferrin intensity at 10 min in each condition in **C**, and immediately after the pulse in each condition in **D**. **E,** Representative maximum intensity projections of fluorescence confocal images of iCE-BECs with ApoE3 or ApoE4 genetic variant after immunostaining for transferrin receptor (TfR). Cells are pseudo-colored showing TfR in green, Phalloidin in magenta and DAPI-stained nuclei in blue. Scale bar, 20 µm. **F,** Quantification of TfR integrated intensity in Phalloidin-positive area. Intensity values for each experiment were normalized to ApoE3 conditions. Graph shows boxplots with interquartile ranges and median. Lines show the 5th and 95th percentiles, data come from n = 3 independent differentiations with 30 images corresponding to approximately 200 cells per condition per experiment. Differences in the TfR intensity between ApoE genetic variants are statistically significant as evaluated by Student’s t test. ***, p <0.001. **G,** Representative immunoblot detecting TfR with GAPDH as a loading control. Data come from three independent differentiations (Diff). **H,** Quantification of relative TfR mRNA expression by quantitative PCR. Graph shows mean ± SD of n = 3 independent differentiations with 3 technical replicates per experiment. Points represent independent differentiations. Differences in TfR mRNA expression are significantly different between ApoE genetic variant as evaluated by Student’s t test. **, p < 0.01.

### ApoE4 alters iron metabolism homeostasis in iCE-BECs

TfR expression is regulated by iron responsive proteins that sense iron levels in the cytoplasm ^51^. In iron-deficient cells, iron responsive proteins upregulate TfR1 mRNA by binding to its 3’UTR thus preventing its degradation ^52^. We therefore tested whether ApoE4 iCE-BECs had lower cytosolic iron levels. To measure the intracellular labile iron pool, we used FerroOrange, a fluorescent probe that specifically detects labile iron (II) ions (Fe2^+^) in live cells. To verify the specificity of the fluorescent probe, we measured the effects of an iron donor (Ferrous ammonium sulfate, FAS) or an iron chelator (pyridoxal isonicotinoyl hydrazone, PIH) and observed increase and decrease of fluorescent intensity, respectively (Figure S7A). With this method, we found that ApoE4 iCE-BECs had a reduced labile iron pool compared to ApoE3 cells (Figure 6A, B). We confirmed this result with an orthogonal method that measures the intracellular iron pool with the metal-sensitive calcein-acetoxymethyl ester (Calcein-AM) ^53^. Briefly, calcein-AM fluorescence is quenched upon binding to iron and unquenched upon treatment with the iron chelator pyridoxal isonicotinoyl hydrazone (PIH). This method confirmed a reduction in the labile iron pool in ApoE4 compared to ApoE3 iCE-BECs (Figure S7B-D).

**Figure 6.**
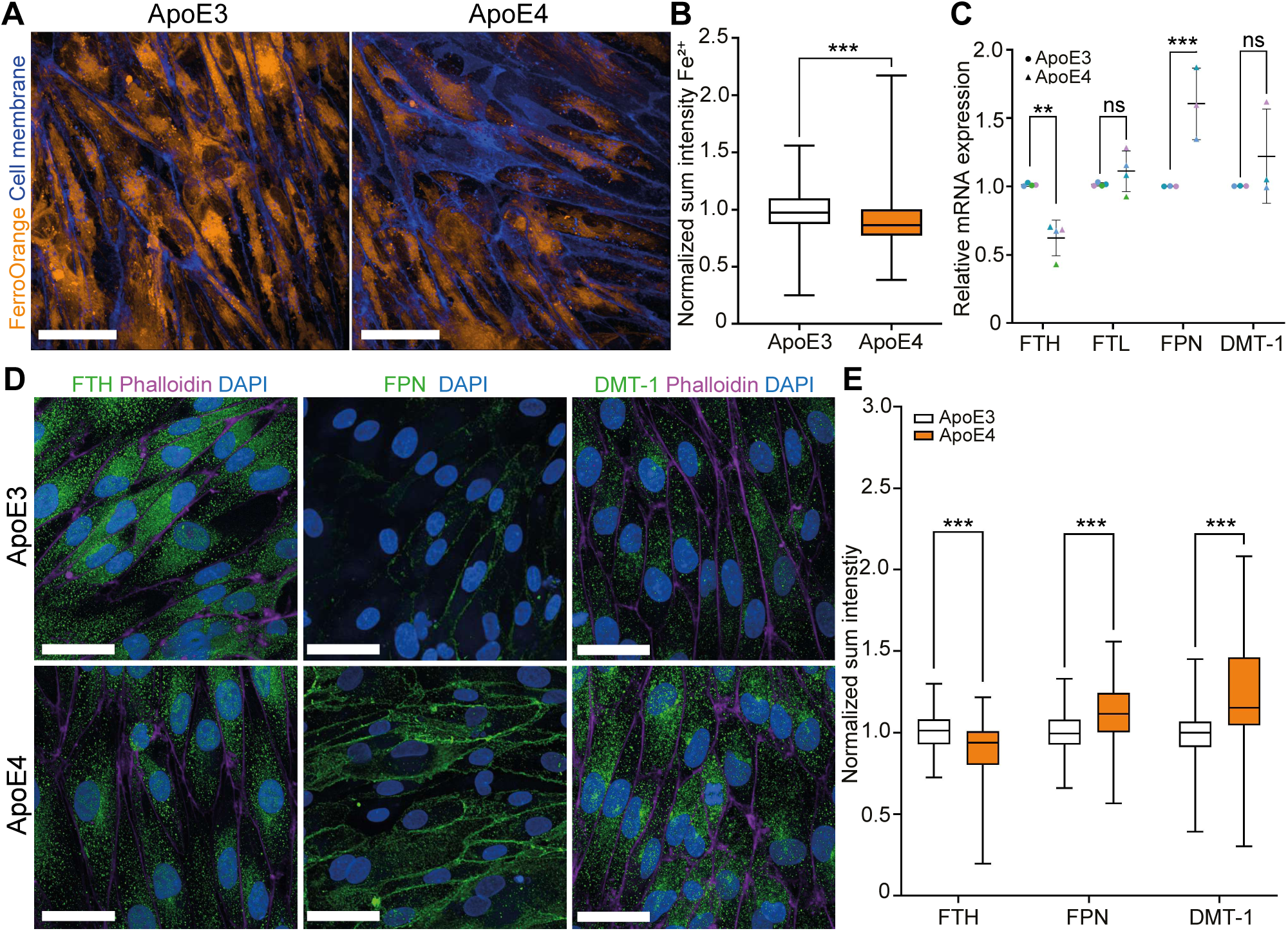
ApoE4 reduces intracellular iron levels and alters iron transport pathways in iCE-BECs. **A,** Representative maximum intensity projections of fluorescence confocal images of iCE-BECs with ApoE3 or ApoE4 genetic variant after labeling labile iron (II) ions (Fe^2+^) with FerroOrange. Cells are pseudo-colored showing FerroOrange in orange, cell area in blue. Scale bar, 50 µm. **B,** Quantification of sum intensity of FerroOrange intensity within cell area. Intensity values were normalized to data from ApoE3 iCE-BECs per experiment. Graph shows boxplots with interquartile ranges and median. Lines show the 5th and 95th percentiles, data come from n = 3 independent differentiations with 40 images per experiment. Differences in the FerroOrange intensity between ApoE genetic variants are statistically significant as evaluated by the Mann-Whitney-U test. ***, p < 0.001. **C,** Quantification of relative mRNA expression by quantitative PCR of Ferritin heavy chain (FTH), Ferritin light chain (FTL), Ferroportin (FPN) and Divalent-metal transporter 1 (DMT-1). Graph shows mean ± SD of n = 3 independent differentiations with 3 technical replicates per experiment. Points represent independent differentiations. Differences in mRNA expression for each gene were statistically evaluated using Two-Way ANOVA and Sidak multiple comparisons. ns, not statistically significant; **, p<0.01; ***, p<0.001. **D,** Representative maximum intensity projections of fluorescence confocal images of iCE-BECs with ApoE3 or ApoE4 genetic variant after immunostaining for FTH, FPN or DMT-1 (green) and actin (phalloidin, magenta). DAPI-stained nuclei are shown in blue. Scale bar, 100 µm. **E,** Quantification of FTH, FPN or DMT-1 sum intensity in Phalloidin^+^ area. Intensity values were normalized to data from ApoE3 iCE-BECs for each experiment. Graph shows boxplots with interquartile ranges and median. Lines show the 5th and 95th percentiles, data n = 3 independent differentiations with 50 images per experiment. Differences in protein expression were statistically evaluated using Two-Way ANOVA with Sidak multiple comparisons. ***, p<0.001.

The changes we observed in the iron labile pool in iCE-BECs suggest that the effect of ApoE4 is not restricted specifically to TfR1 but could broadly impact iron metabolism. Therefore, we evaluated proteins regulated by iron responsive elements involved in different stages of the iron transport pathway: Ferritin (Ferritin heavy chain, FTH; Ferritin light chain, FTL; both iron storage), Ferroportin (FPN, iron export) and divalent metal transporter 1 (DMT-1, iron import). It is well documented that low cytosolic iron downregulates FPN and FTH/FTL while upregulating DMT-1 ^52,54^. We found that Ferritin mRNA and protein expression was downregulated whereas DMT-1 was upregulated in ApoE4 compared to ApoE3 iCE-BECs (Figure 6C, E). Interestingly, DMT-1 signal was accumulated in vesicular structures and a perinuclear compartment (Figure 6D). Unexpectedly, FPN mRNA and protein expression was increased and was relocalized to the plasma membrane (Figure 6C-E). Altogether, our data show that ApoE4 alters iron cytoplasmic levels and leads to expression changes across the iron transport pathway in iCE-BECs.

In conclusion, ApoE4 iCE-BECs exhibited altered spatiotemporal organization of endosomes and disrupted iron homeostasis, characterized by lower intracellular iron levels and altered expression of iron-related genes, including upregulation of TfR1. These findings highlight the importance of the ApoE4 gene variant in modulating intracellular trafficking and iron metabolism within brain endothelial cells.

## Discussion

Recent work highlighted the importance of developing and optimizing iPSC-based protocols to improve brain endothelial identity in order to investigate specific physiological mechanisms ^26,55^. Here, we describe an efficient and high-yield protocol to generate iCE-BECs that shows 1) an endothelial transcriptomic signature, 2) improved barrier properties and 3) recapitulates receptor-mediated transcytosis. The iCE-BEC protocol shares similarities with the recently described cARLA differentiation method ^56^ and thus confirms the importance of Wnt/cAMP activation and TGF-β inhibition to robustly induce brain endothelial identity and barrier properties of endothelial cells *in vitro* ^57^. However, when compared with the cARLA differentiation method, iCE-BECs show increased expression of ZIC3 and FOXF2, two transcription factors specifically enriched in brain endothelial cells *in vivo* and important to induce barrier properties in endothelial cells ^58^. A recent human model for transcytosis using cells with mixed epithelial identity observed antibody transcytosis at concentrations at least 10-fold higher than previously reported ^38^. In contrast, iCE-BECs showed the same extent of transport using similar concentrations as reported by a BBB organoid model ^41^. We therefore consider iCE-BECs a suitable model to investigate receptor-mediated transcytosis across the BBB and, more specifically, how disease-related mechanisms impact BBB transport.

The ApoE4 genetic variant is the major risk factor for sporadic AD ^20,42,43^ heavily affecting the onset of the disease ^59^. Extensive evidence documents the impact of ApoE4 on BBB permeability and function ^16–21^. In this manuscript, we demonstrate for the first time how ApoE4 expression in brain endothelial cells leads to both disruption of sorting tubule formation and altered iron homeostasis. This finding expands the role of ApoE4 at the BBB beyond paracellular permeability to include the cell-autonomous regulation of intracellular transport. Previous studies using human tissue ^22,60^ or mouse models ^60,61^ did not find upregulation of TfR1 at the BBB in AD. However, these studies were not designed to evaluate the specific role of ApoE4 on BBB phenotypes. In contrast, an ApoE4 knock-in mouse line showed an increased in TfR1 expression in brain endothelial cells ^23^, similarly to the effect that we see in our in vitro model. Our data suggest that TfR1 upregulation is a consequence of lower cytoplasmic iron levels in ApoE4 cells. We propose a model that could explain how changes to the endosomal network observed in ApoE4 BECs lead to lower cytosolic iron levels (Figure 7). First, enlarged endosomes and reduced sorting tubule formation could alter endosome maturation/acidification ^62^. In agreement with this hypothesis, iPSC-derived astrocytes with the ApoE4 gene variant also showed enlarged early endosomes ^63^ and altered endolysosomal pH ^64^. Second, changes in endosomal pH could reduce dissociation of ferric iron from transferrin. Third, sustained ferric iron binding to transferrin would prevent DMT-1-mediated transport into the cytosol. Together, these events would lead to an iron-depleted cytosol and trigger expression changes iron-related genes, including TfR1 (Figure 7).

**Figure 7.**
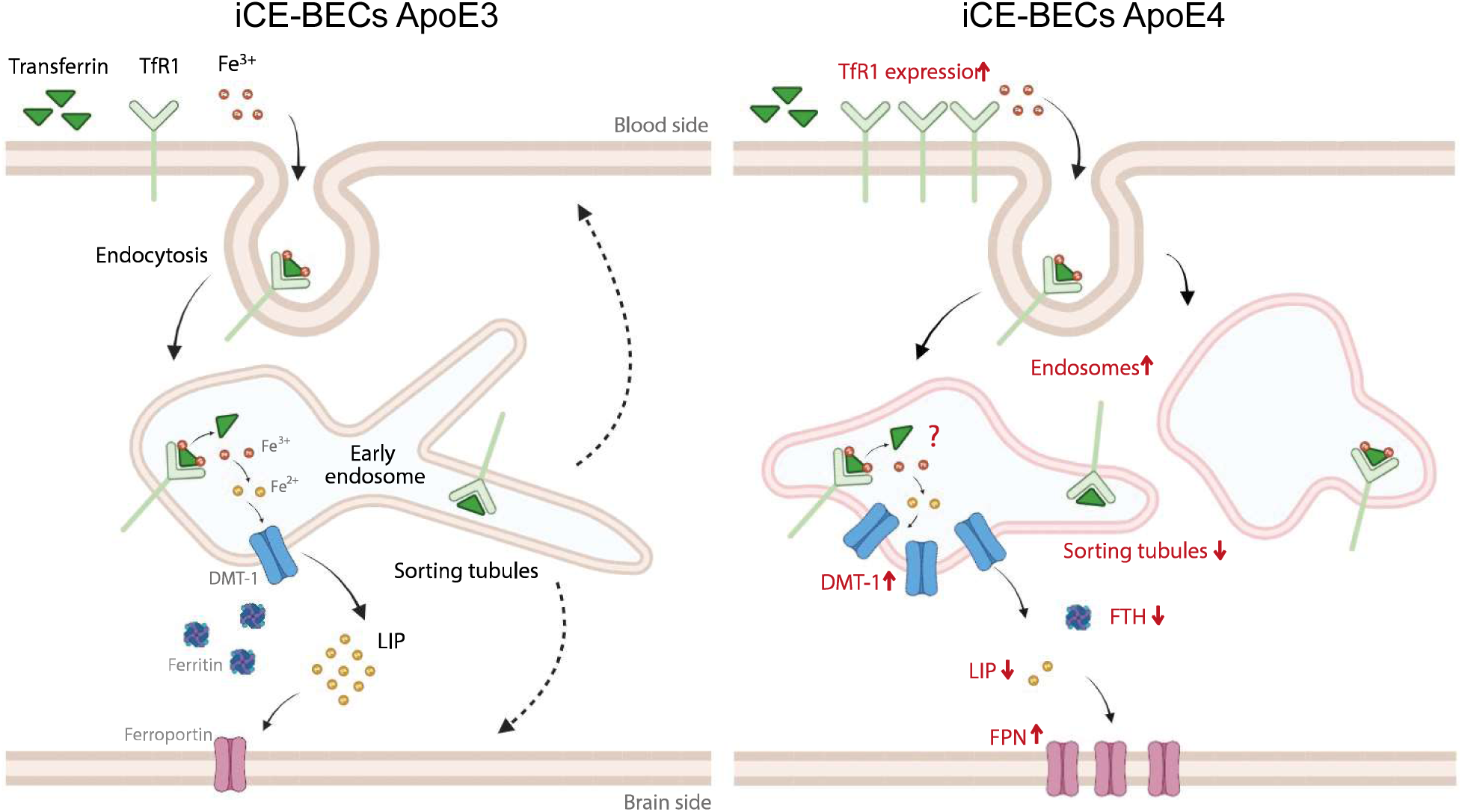
ApoE4 impacts iron homeostasis in iCE-BECs. Schematic summarizing intracellular transport of transferrin and iron in BECs with ApoE3 genetic variant and the key changes occurring in ApoE4 iCE-BECs: a) early endosome enlargement and reduced sorting tubule biogenesis. b) Reduced intracellular labile iron pool (LIP), potentially caused by defects in endosomal maturation. c) Changes in expression of proteins regulated by iron-responsive elements, including Divalent metal transporter 1 (DMT-1), Transferrin receptor TfR (both increased) and Ferritin (FTH, also reduced). Increased Ferroportin expression could be caused by additional feedback mechanisms. d) Increased transferrin uptake driven by higher TfR1 expression.

How ApoE4 leads to changes in early endosome organization still remains to be clarified. Our live-cell imaging data suggest that endosome enlargement is caused by reduced sorting tubule biogenesis, which is regulated by the recruitment and assembly of sorting complexes ^65^. Transcriptomic analysis performed on brains of ApoE4 mice revealed a significant upregulation of genes involved in the regulation of endosomal–lysosomal pathway, including *Rab5b, Rab7, Snx3, Snx15, Vps4a, Vps24, Vps29,* and suggested an ApoE4-specific trafficking and sorting dysregulation ^49^. Similarly, brain endothelial cells purified from ApoE4 mice, showed alterations in the expression of genes involved in subcellular vesicle trafficking and clathrin-mediated transport ^23^. Alternatively, changes to lipid metabolism triggered by ApoE4 ^66–70^ could change the endosomal membrane composition and thus impact the recruitment of lipid-dependent sorting adaptors. Follow-up studies with iCE-BECs could help to dissect the specific molecular mechanisms that trigger the changes we observed in the endosomal network. Overall, our findings on the influence of ApoE4 on endosomes and TfR1 expression at the BBB will be instrumental in refining strategies that leverage intracellular transport mechanisms for the delivery of therapeutic antibodies to the brain.

### Limitation of the study

While ApoE is expressed in endothelial cells, this is not the primary source of ApoE in the brain. It was reported that astrocyte-produced ApoE4 can lead to BBB impairment ^71^, and it could also impact the changes we described on iCE-BECs. Our experimental setup utilizes a monoculture system that primarily investigates cell-autonomous mechanisms. Consequently, we are unable to assess the impact of interactions within a more complex, multicellular environment or the effect on barrier properties, given the lack of pericytes and astrocytes. Additionally, the translatability of the phenotypes described in iCE-BECs still needs to be confirmed in patients.

## Experimental procedures

### Resource availability

Requests for further information or more detailed protocols should be directed to and will be fulfilled by the corresponding author, Roberto Villasenor Solorio.

### Material availability

This study did not generate new unique reagents.

### Data and code availability

Data will be shared with the research community upon request. No code or standardized datasets were generated.

### hiPSC lines

For protocol comparison and model establishment, experiments were performed differentiating endothelial cells from hiPS_SFC086_03_03 line. hiPS_SFC086_03_03 were established by reprogramming skin fibroblasts with Sendai-Virus (cytotune v II kit) at STEMBancc from a female donor. Karyotype analysis was performed by WiCell and no abnormalities were detected. Additionally, to assess reproducibility of the protocol, BIONi010-C13 (European Bank for induced pluripotent Stem Cells, ebisc.org), Alstem iPS26 (see below), and BIONi037-A (see below) were used. For ApoE experiments, Bioni037-A (16423, homozygous ApoE3 gene variant) and Bioni037-A4 (I40-53, homozygous ApoE4 gene variant) were used. The parental homozygous ApoE3 line was generated from the fibroblasts of a healthy individual with homozygous ApoE3 genetic variant ^46^.The isogenic ApoE4 homozygous line was derived from the parental line by editing of the ApoE allele from T/T to C/C at rs429358, which changes aa from Cys112 to Arg112; the genetic variant of base position described by rs7412 in both parent and subclone is C/C, which is an Arg. Together these define the ApoE4/E4 genetic variant in this subclone. Editing was confirmed and cells were characterized (sequencing, expression of pluripotency markers, morphology) by the European Bank for induced pluripotent Stem Cells (ebisc.org). For validation of the phenotype, Alstem line iPS16 (homozygous ApoE4 gene variant) and iPS26 (homozygous ApoE3 gene variant) were used. The parental Alstem line iPS16 was reprogrammed from one single iPSC clone of human bone marrow CD34-positive mononuclear cells. The isogenic control line carrying ApoE3 (iPS26, Alstem) was derived from the parental Alstem line by changing Arg112 to Cys112 in the ApoE allele. Editing was confirmed and cells were characterized (sequencing, expression of pluripotency markers, morphology) by Alstem (alstembio.com). All cell lines were tested for mycoplasma contamination.

### hiPSC differentiation into iCE-BECs

hiPSC were maintained in culture under standard culturing conditions using plates coated with Geltrex (A1413301, ThermoFisher), mTeSR Plus Media (100-0276, Stemcell) and passaging done with Gentle Cell Dissociation Reagent (100-0485, Stemcell). To differentiate iCE-BECs, 2 million hiPSCs were seeded in a 10 cm dish coated with Geltrex in 10 mL of mTeSR Plus Media supplemented with ROCK inhibitor Y-27632 10 uM (SCM075, EMD Millipore). 24 h later the media was replaced with mesodermal induction media composed by DMEM/F12 (31331-028, Gibco) and Neurobasal medium (21103-049, LifeTechnologies) 1:1, 2-Mercaptoethanol (31350-10, ThermoFisher), B27 (17504044, Gibco), N2 (17502048, Gibco) supplemented with fresh CHIR-99021 8 μM (13122, Cayman) and BMP4 25ng/mL (120-05ET, Peprotech). At day 4 and 5 media was replaced with Endothelial induction media composed by StemPro-34 SFM Media (10639011, Life Technologies) with StemPro-34 supplement, Glutamax (35050061, Gibco), Penicillin-Streptomycin (15070063, Gibco) freshly supplemented with VEGF 50ng/mL (293-VE-010, R&D) and Forskolin 2uM (ab120058, abcam). At day 6 in culture, cells were replated in 10 cm dishes (1.2 million per dish). Dishes were coated with Vitronectin 2.5 µg/ml (SRP3186, Sigma) and Fibronectin 7.5 ug/ml (F0895, Sigma) and cells were kept in BBB Identity Acquisition media consisting of Vasculife VEGF Endothelial Medium (LL-0003, Lifeline Cell technology) supplemented with iCell Endothelial cells medium supplement (M1019, Fuji cell dynamics) instead of the FBS included in the kit and just 10 mL of L-Glutamine LifeFactor instead of the 25 mL included in the kit. The media was freshly supplemented with CHIR-99021 4 µM (13122, Cayman), SB-431542 5 μM (72234, Stemcell), cAMP 50 nM (ab120424, abcam). Cells were cultured in these conditions up to day 11 changing media every 2 or 3 days. At day 11 in culture, CD31-positive cells were MACS sorted according to the manufacturer’s protocol using CD31 MicroBead Kit from MACS Miltenyi Biotec (130-091-935, Miltenyi Biotec). After MACS, cells were either frozen in liquid nitrogen or replated in the same conditions with BBB Identity Acquisition media. For protocol comparison (schematic in Figure 1A), at day 6, cells were replated in just Fibronectin coated dishes in Vasculife VEGF Endothelial Medium with the modifications aforementioned and without CHIR-99021, SB-431542 and cAMP (iEC and iEC-rep). For For iEC-rep, media was supplemented with RepSox 10 µM (73794, Stemcell) 48 h before performing the analysis. For all experiments performed with iCE-BEC, plates and dishes were coated with Vitronectin and Fibronectin prior to seeding of the cells and cells were cultured in BBB Identity Acquisition media. Experiments have been carried out on day 14 (three days after MACS sorting), if not indicated differently.

### Single cell RNAseq sequencing and analysis

300,000 cells differentiated to iECs, iEC-rep, iCE-BECs (see figure 1A) were plated in 6 well plates at day 11 and grown at confluence to day 14. At day 14, cells were detached and resuspended into a single cell suspension concentrated 1×10^6^ cells per mL. Cells were then processed following the Chromium Next GEM Single Cell 3’ v3.1 protocol for GEM generation and barcoding followed by gene expression library construction according to the manufacturer’s instructions. Targeted cell recovery of 10’000 per sample was performed. Dual indexed libraries were sequenced on the Novaseq 6000 with a target sequencing depth of 50’000 single reads per cell. Raw reads were processed with Cellranger software (version 7.1.0) and aligned to hg38 reference transcriptome.Count matrices generated by Cell Ranger were corrected for ambient RNA using CellBender 0.2.0 with the expected cells parsed from the Cell Ranger web summary. The number of total droplets to include was set to 25000, the false positive rate set to 0.01 and the algorithm was trained for 150 epochs. CellBender corrected count matrices were then concatenated and further processed with scanpy (v1.9.3). Barcodes were filtered for observations that CellBender had assigned a latent cell probability greater than 0.5. Furthermore, we excluded low quality cells and potential duplicates by retaining cells with percentage of mitochondrial counts < 5% and with a number of genes between 200 and 2500. Genes detected in less than 30 cells were removed from downstream analysis. Counts were normalized by library size and log1p transformed before determining highly variable genes and regressing out the effect of library size and percentage of mitochondrial counts. Top 10 principal components were used as input for the neighborhood graph which in turn was used to create the UMAP representation.

We utilized the metaanalysis published by Liu and colleagues ^26^ to generate genesets associated with an endothelial transcriptomic signature (Positive PC1 loading) or an epithelial transcriptomic identity (Negative PC1 loading). In detail, we extracted the top 500 genes with strongest positive and 500 genes with strongest negative contribution to PC1 from sheet “Fig.1C PC1 Loading Genes” from dataset S02 provided as supporting information to their publication (see Supplementary Table 1). We subset their genes to those expressed in our data (392 for positive PC1 loading, 193 for negative PC1 loading) and visualized the score for the two genesets through the Seurat (v5.0.1) function AddModuleScore.

### Bulk RNA sequencing

300,000 cells differentiated to iEC, iEC-rep or iCE-BECs (see figure 1A) were plated at day 11 and grown at confluence to day 14. Differentiation protocols were tested in triplicate, yielding a total of 12 samples (three independent differentiations per condition) for sequencing. At day 14, cells were collected and pellets were snap frozen. RNA was extracted and preparation of Illumina stranded TruSeq RNA libraries, including poly(A) enrichment, was performed (2*100 bp paired-end reads). Samples were sequenced on the Illumina NovaSeq and NextSeq by Microsynth. Base calling was conducted using the BCL to FASTQ file converter bcl2fastq2 version 2.20.0 (Illumina). Quality assessment of FASTQ files was performed with FastQC version 0.12.1 (Andrews et al. 2010). Paired-end reads were aligned to the human genome (build “hg38”) using the STAR read aligner version 2.7.11b with default mapping parameters (Dobin et al. 2013). Alignment metrics were determined using Picard version 3.1.1 (Broad Institute). Quality of read sequences and alignments was assessed with MultiQC version 1.21 ^72^. The number of reads mapped to all RefSeq transcript variants of a gene were combined into a single count value (i.e., read count) assuming a reverse-stranded library, using featureCounts version 2.0.6 ^73^. Read counts for 43,294 RefSeq transcripts were generated for all 12 samples. All samples passed quality control checks. Read count normalization, principal component analysis (PCA) and gene ontology (GO) term enrichment were subsequently conducted in R (version 4.3.0). Read count normalization was performed using the edgeR package (version 4.0.5; ^74^). Library size and composition were adjusted for using Trimmed mean of M (TMM)-normalized CPM values. Only samples with >=1 CPM and >10 read counts in at least 3 samples were retained. Following gene filtering, a total of 13,731 of 43,294 transcripts were identified as expressed (32%). The PCA was computed based on the top 500 most variable expressed genes using TMM-normalized log2(CPM+1) values. Eigenvalues were extracted and the cumulative percentage of each principal component’s (PC) variance explained. PC1 contributed most to sample variance (78.5%), with PC2-4 individually contributing <7% each. GO term enrichment was performed on the top 100 loading genes for PC1 using the clusterProfiler package in R (version 4.10.0). Multiple testing correction was performed using the Benjamini-Hochberg method.

### Whole cell proteomics

To benchmark iCE-BECs, primary brain endothelial cells (HBMVEC, n=6 from three different batches), immortalized endothelial cells (HCMEC/D3, n=3 different passages), and induced ECM-supported brain endothelial cells (iCE-BECs, n=4 differentiations) were analyzed. HCMEC/D3 (SCC066, Merck) cells were cultured in EGM-2 Endothelial Cell Growth Medium-2 BulletKit (CC-3162, Lonza) for three sequential passages and collected when at confluence. Three independent batches of HBMVECs were purchased from AngioProteomie (cAP-0002, AngioProteomie) and cultured in EGM-2 Endothelial Cell Growth Medium-2 BulletKit (CC-3162, Lonza) in flasks coated with quick coating solution (cAP-01, Angioproteomie). Each cell batch was cultured for two subsequent passages and collected when at confluence. iCE-BECs were differentiated as previously described and collected at day 14 in culture. For whole proteomics analysis, culture media were removed and cells were washed with phosphate buffered saline (PBS). Cells were detached using trypsin 0.25%-EDTA (ThermoFisher, 25200056) for HCMEC/D3 or TrypLE^TM^ (12563011, ThermoFisher) for HBMVECs and iCE-BECs, resuspended in culture medium and centrifuged at 180 g at 4°C for 5 min. Cell pellets were then washed with cold PBS and centrifuged. This washing step was repeated again once before snap freezing the cell pellets on dry ice for about 15 min. Cell pellets were then stored at -80°C.

Samples were reduced, alkylated, digested with trypsin and peptides purified using the PreOmics iST kit according to the supplier’s specifications. Peptide samples were resuspended in 2% (v/v) acetonitrile and 0.5% (v/v) formic acid solution and 1ug of peptides were analyzed by liquid chromatography (nano capillary system, EASY-nLC™ 1200 system, Thermo Scientific) on a C18 reverse-phase nano-high-performance liquid chromatography column connected to a mass spectrometer (Orbitrap Exploris^TM^ 480, Thermo Scientific) via electrospray ionization. The DIA method consisted of one full range MS1 from 340 to 1210 m/z at 120k resolution, with a custom AGC target and 20ms max injection time. Then 28 DIA segments were acquired at 15k resolution with a standard AGC target and 20 ms max injection time. HCD fragmentation was set to normalized collision energy optimized for each segment. The spectra were recorded in profile mode. The default charge state for the MS2 was set to 3. Raw files have been processed with Spectronaut 18, with experiment settings based on BGS Default SNE for a DIA library free search, using global imputation to deal with missing values. Default settings included peptide and protein level false discovery rate control at 1 %. Measurements were normalized separately using local regression normalization. The mass spectrometric data were analyzed using Pulsar search engine as implemented in Spectronaut software, the false discovery rate on peptide and protein level was set to 1 %. A human UniProt fasta database (Homo Sapiens, 2022 07 01) was used for the search engine, allowing for 2 missed cleavages and variable modifications (N term acetylation and methionine oxidation).

Distributions of both raw and normalized data at the protein level were assessed to evaluate sample consistency. Principal Component Analysis (PCA) was conducted to reveal the overall data structure and to identify potential experimental artifacts. Outlier detection was implemented using the Mahalanobis distance, calculated from the first three principal components. No outliers were removed. The differential abundance analysis was performed to identify proteins with significant changes in expression levels between the different conditions. For this statistical analysis, we utilized the R package “limma”. Unlike standard t-tests, which compare proteins individually, the “limma”; approach accounts for the overall variance observed across all proteins. This typically results in adjusted P-values, especially at the tails of the distribution, and is particularly well-suited for studies with small sample sizes.

### FITC-dextran Permeability in Transwell

50,000 cells were plated at day 11 on the insert of transwell chambers (734-4072, Avantor). Media was changed at day 12 and FITC-dextran permeability experiments were run at day 14 in culture. Briefly, the inserts were moved to a new Receiver Tray supplemented with 600 µL of fresh media. A FITC-dextran (3.3 kDa, D3305; 40 kDa, D1845; 70 kDa D1822, all purchased from ThermoFisher) dilution of 50 µg/ml was added in the insert compartment and plates were incubated for 30 minutes at 37 °C. After incubation, 100 μL of the media from each Receiver Tray were transferred to wells of a black 96-well opaque plate (PBK96G-1.5-F, MatTek USA) for fluorescence measurement. Fluorescence was read at 485 nm and 535 nm excitation and emission, respectively and apparent permeability (P_app_) calculated using the formula:

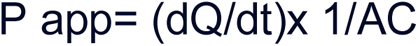

where dQ/dt is change in concentration / change in time, A is the growth area in the insert and C is the initial concentration in the insert chamber.

### Histology of transwell

iCE-BECs were seeded on a transwell as described above and fixed at day 14 in 4% PFA for 30 min, subsequently washed with PBS for three times. Cells including the transwell mesh were embedded in 2% Agarose (V3121, Promega), dehydrated overnight (TissueTek VIP5, Sakura) and ultimately embedded vertically in paraffin using an embedding console (Tissue-Tek® TEC™5, Sakura). Sagittal microtome sections at 4 um were prepared on Superforst Plus glass slides (J1800AMNZ, ThermoFisher) and an automated Haematoxylin & Eosin staining was performed (Ventana HE600, Roche).

### High content live imaging of Brainshuttle™ transcytosis and permeability in microphysiological system

To test the barrier function of iCE-BECs in a microphysiological system, cells were seeded in an OrganoPlate 2-lane (9605-400-B, MIMETAS) according to the manufacturer’s protocol ^39^. Briefly, one day before the cell seeding, ECM (3447-020-01, Cultrex 3D Collagen I, R&D Systems in 1M Hepes, Gibco and 37 mg/mL NaHCO_3_) was prepared. Per tube, 35.000 cells were seeded and supplied with complete BEC media. After 72 h, media was replaced with starvation media (1:5 complete BEC media : basal media without supplements) for basal conditions or additionally supplemented with 0.2 µg/mL VEGF (293-VE-010, R&D). After 24 h incubation, differently sized FITC-dextrans (3.3 kDa, D3305; 40 kDa, D1845; 70 kDa D1822, all purchased from ThermoFisher) were applied at 10 µg/mL and immediately imaged using Opera Phenix High Content Imaging System (PerkinElmer) at 5× magnification, every five minutes for two hours. The ratio between mean intensity of FITC-dextran in the cell channel and the gel channel was calculated for each perfusable tube at each time point.

To calculate the apparent permeability, the slope of the linear regression was multiplied by the volume of the gel (0.0004136 cm^3^) and subsequently divided by the surface area (0.01218153 cm^2^) adapted from ^75^ according to the manufacture’s protocol.

To assess transcytosis of Brainshuttle^TM^ molecules across iCE-BECs, cells were seeded in an OrganoPlate 2-lane (9605-400-B, MIMETAS) as described above. After starvation for 24 h, fluorescently labeled Brainshuttle^TM^ constructs or non-targeting IgGs were applied at 200 nM, and immediately imaged using Opera Phenix High Content Imaging System (PerkinElmer) at 5× magnification, every 15 min for twelve hours. The ratio between mean intensity of fluorescently labeled Brainshuttle^TM^ or non-targeting IgG in the donor cell channel and the gel channel was calculated for each perfusable tube at each time point. To assess total transcytosis of Brainshuttle^TM^, the slope of the linear regression was calculated within the time interval 0 to 12 h. Before each experiment, channels with matrix overflow or incomplete filling of matrix channel were excluded from the analysis.

### Transferrin kinetics (uptake and pulse-chase)

To assess the uptake amount and rate of transferrin (pulse assay) and recycling rate of transferrin (pulse-chase assay), 25.000 cells were seeded in a 96-well glass bottom imaging plate (PBK96G-1.5-F, MatTek USA) in BBB Identity Acquisition media on day 11. On day 14, cells were washed once with PBS and incubated with the assay medium (EGM, cAP-02, Angioproteomie) containing 1% Bovine serum albumin for at least 10 min. Labeled transferrin (T13342, ThermoFisher) at 25 µg/mL was then applied between 2 and 45 min to assess the time course of transferrin uptake. To assess the recycling rate, cells were treated with labeled transferrin (T13342, ThermoFisher) at 25 µg/mL for 20 min, followed by application of 10-fold higher concentration of unlabeled Holo-Transferrin (T0665, 250 µg/mL) for different time points (0 min - 60 min). Cells were fixed in 4% PFA for 20 min and washed three times with PBS before counterstaining with Phalloidin-Atto-647 (65906, Sigma) and DAPI (D9542, Sigma). Images were acquired with Opera Phenix High Content Imaging System (PerkinElmer) using the 40× water long WD confocal objective, Binning 2×2, Camera ROI 2160×2160, 20 planes à 1 μm. Per time point, minimum 50 images were acquired.

### Live imaging of sorting tubule biogenesis

To assess the formation of sorting tubules, 15.000 cells were seeded in a 96-well glass bottom imaging plate (PBK96G-1.5-F, MatTek USA) in BBB Identity Acquisition media 48 h before the experiment. After washing with PBS, cells were incubated with 25 µg/ml of fluorescently labeled Transferrin (T13342, ThermoFisher) for at least three hours at 37°C 5% CO_2_. After washing with BBB Identity acquisition media, cells were imaged with a DMI-8 TIRF microscope (Leica Microsystems) equipped with a stage incubator controlling at 37 °C and 5% CO_2_. Full frame 1024 x 1024 images were acquired with HCX PL APO 100X/1.4 NA oil objectives with a resolution of 0.288 µm, two z-stacks with 1 µm step size. Single color images were acquired with a 488 nm TIRF laser for a final rate of 2 frames per second for 1 min each. The number of tubules per cell occurring within 1 min acquisition was quantified manually in the maximum intensity projection of the z-sections for each movie by two independent blinded raters.

### Immunostainings

After fixation of the cells with 4% PFA (15710, Electron Microscopy Science), cells were quickly washed with PBS for three times, before permeabilized with 4% gelatin (G7041, Sigma) + 0.1% Saponin (84510, Sigma) in PBS for 10 min at RT. Primary antibodies were incubated overnight at 4 °C (see Supplementary Table 1). After washing three times with PBS, fluorescently labeled secondary antibodies and Phalloidin-Atto-647 (65906, Sigma) were applied for one hour at room temperature (RT, see Supplementary Table 2). Nuclei were counterstained with DAPI (Sigma, D9542) for 10 min at RT. For representative images, confocal images at SP-8 using 63× or 25× objective or a DMI-8 TIRF microscope at 20× magnification (Leica Microsystems) were acquired. For quantification of iron-related proteins, Opera Phenix High Content Imaging System (PerkinElmer) was used at 40× magnification.

**Table 1.** Primary antibodies used for immunostainings.

| Target | Concentration | Catalog No. | Vendor |
| --- | --- | --- | --- |
| Early Endosome Antigen 1 (EEA1) | 1:200 | 3288S | Cell signaling |
| Transferrin Receptor (TfR) | 1:200 | 13-6800 | ThermoFisher |
| VE-Cadherin | 1:200 | 2500S | Cell signaling |
| PECAM1 | 1:200 | NB600-562 | Novus Biologicals |
| Ferritin Heavy chain (FTH) | 1:200 | ab65080 | Abcam |
| Zonula occludens-1 | 1:200 | 33-9100 | ThermoFisher |
| (ZO-1) |  |  |  |
| Divalent metal transporter 1 (DMT-1) | 1:200 | ab55735 | Abcam |
| Ferroportin (FPN) | 1:200 | HPA065634 | Sigma |
| Claudin-5 | 1:200 | 352588 | ThermoFisher |

**Table 2.**
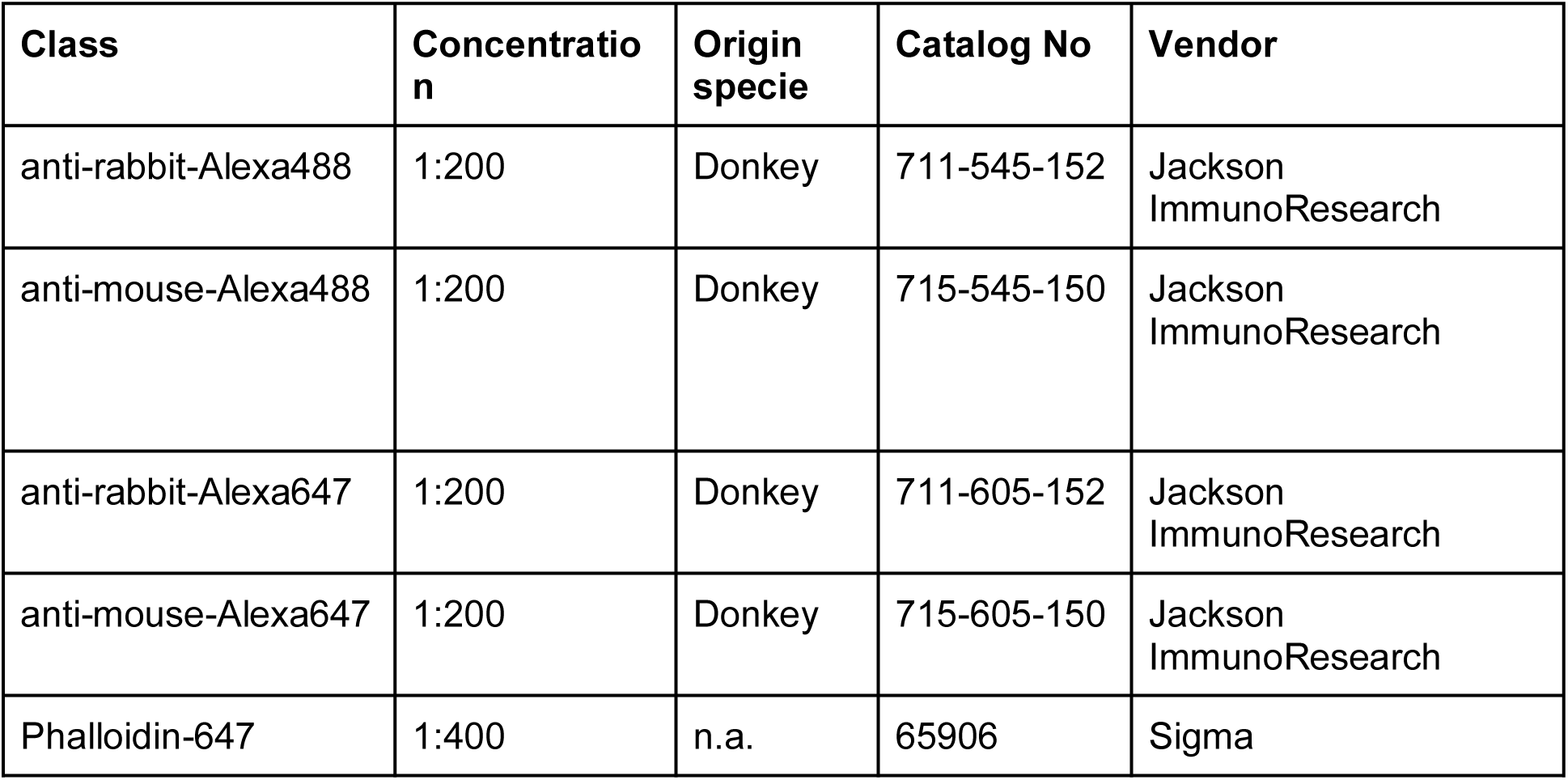
Secondary antibodies and fluorescent probes for immunostainings.

| Class | Concentration | Origin specie | Catalog No | Vendor |
| --- | --- | --- | --- | --- |
| anti-rabbit-Alexa488 | 1:200 | Donkey | 711-545-152 | Jackson ImmunoResearch |
| anti-mouse-Alexa488 | 1:200 | Donkey | 715-545-150 | Jackson ImmunoResearch |
| anti-rabbit-Alexa647 | 1:200 | Donkey | 711-605-152 | Jackson ImmunoResearch |
| anti-mouse-Alexa647 | 1:200 | Donkey | 715-605-150 | Jackson ImmunoResearch |
| Phalloidin-647 | 1:400 | n.a. | 65906 | Sigma |

### High content imaging and image analysis for transferrin kinetics and immunostainings

The Perkin Elmer’s Harmony high-content analysis software 5.1 (HH17000001) was used to set up the plate dimensions enabling the fast and efficient imaging with the Opera Phenix High Content Imaging System (PerkinElmer). For transferrin kinetics and immunostainings for iron-related protein experiments, we used the 40× long WD confocal objective. Per well, 50 images were acquired, 20 planes per image with a section thickness of 1 µm per plane. Appropriate channels were selected (DAPI, Alexa-488, Alexa-647) and exposure time as well as focus height was set accordingly and kept the same between wells. Perkin Elmer’s Harmony high-content analysis software 5.1 was used to analyze the images. In brief, basic flat field correction and maximum projection was applied. Cell area was determined by a binary threshold mask using the Phalloidin signal. Sum intensity of the respective marker (Transferrin-488, FTH, FPN, DMT-1) was calculated within the cell area and normalized to its area. Linear background reduction was applied. For assessment of the transferrin uptake rate, intensity values were normalized to the 10 min time point, while for transferrin recycling rate, intensity values were normalized to the time point of 20 min pulse, no chase.

### Confocal microscopy and image analysis

Representative images for PECAM1, VE-Cadherin, ZO-1 and Claudin-5 were acquired using a Leica SP-8 HCX PL APO CS at 63× magnification (Leica) or a DMI-8 TIRF microscope at 20× magnification (Leica Microsystems). For quantification of TfR or EEA1 in iCE-BECs, ten images per condition were acquired using the confocal microscope at 63× magnification. Pixel size was adjusted to 100 nm at 1024×1024 frame size. Per image, five z-stacks with a step size of 0.5 µm were used. Pinhole was adjusted to an optical thickness of 1 um and sequential acquisition between stacks was selected. Laser power was adjusted according to the marker expression and not changed between experimental groups. Vesicles containing EEA1 or TfR were detected and mean integrated vesicular intensity within cells (absolute threshold for Phalloidin) was calculated, normalized to the cell area (MotionTracking 8.97, http://motiontracking.mpi-cbg.de/get/) as previously described ^76^.

### Electron microscopy

To assess the ultrastructural morphology of endosomes, cell pellets of iCE-BECs were generated and fixed in 2.5% Glutaraldehyde (pH 7.4) overnight. After lipid fixation with 1% Osmium-Tetroxid for 1 h, samples were dehydrated with ascending ethanols and finally infiltrated with resin by incubating two times with Propylenoxid for 15 min each and Propylenoxid/epon (1:1 ratio) overnight at RT. Samples were transferred to epon blocks and polymerized at 60°C for 60 hours. Ultrathin sections (98 nm) were prepared on 200 mesh copper grids (EMS, Fort Washington, PA, USA) and afterwards contrasted with lead citrate and uranyl acetate. The sections were examined with a Philips CM10 transmission electron microscope equipped with a charge-coupled-device camera (Ultrascan 1000; Gatan) at an acceleration voltage of 80 kV.

### Quantitative PCR

mRNA was extracted from cell cultures using Total RNA Miniprep Kit (T2010, Monarch) following the manufacturer’s instructions. Complementary DNA was synthesized using iScript cDNA Synthesis kit (1708890, BioRad). Quantitative real-time PCR analysis was performed with Lightcycler 480 SYBR Green I Master mix (04887352001, Roche; LightCycler® 96 System, Roche). Ready-to-use primers from Origene for ApoE (HP200028), TFR (HP206788), FTH (HP205786), FTL (HP200131), FPN (HP210988), DMT-1 (HP200584), and GAPDH (HP205798) were used. Primer efficiency was determined by titration of cDNA from iCE-BECs ApoE3; all tested primers had an efficiency between 80%-110%. The cycle threshold (Ct) values were used for all experiments and were first normalized to endogenous control (GAPDH) levels by calculating the ΔCt for each sample. Values were then analyzed relative to control, to generate a ΔΔCt value. Fold change was obtained using the equation, expression fold change=2^−ΔΔCt^.

### Immunoblot

Cells were lysed with RIPA buffer (89900, LifeTech) and incubated at 4°C for 30 min on a rotary shaker. Cells were sonicated for one minute before protein concentration was measured by the bicinchoninic acid assay method (23225, ThermoFisher). Samples were denatured at 95°C for 10 min. Per sample, 10 µg of protein were loaded in 2x Lämmli Buffer containing β-Mercaptoethanol on a Tris-Glycin-Gel (4568096, BioRad). Immunoblots were transferred on membranes (Trans-Blot Turbo Transfer System, BioRad) and subsequently blocked with 5% Milk in Tris-buffered saline with 0.1% Tween® 20 Detergent (TBS-T) for 1h followed by antibody incubation in 5% Bovine Serum Albumin (BSA) in TBS-T overnight (TfR, 13-6800, ThermoFisher, 1:250; ApoE, Ab947, abcam, 1:500). After three times washing in TBS-T, appropriate HRP-conjugated secondary antibodies in 5% BSA in TBS-T were applied (donkey-anti-rabbit-HRP, A16035, ThermoFisher, 1:2000; donkey-anti-goat-HRP, A16005, ThermoFisher, 1:2000) for 1h at RT. Immunoblots were washed three times in TBS-T for five minutes each, before peroxidase (34076, ThermoFisher) was applied. Subsequently, image acquisition was performed with ChemiDoc MP (BioRad).

### Proliferation assay

To assess the proliferation rate between iPSC with ApoE3 and ApoE4 genetic variant, iPSCs were seeded on geltrex (A1413301, ThermoFisher) coated 96-well plate (PBK96G-1.5-F, MatTek USA, 12.000 cells/well) in ROCK inhibitor Y-27632 containing (SCM075, EMD Millipore) mTeSR Plus media (100-0276, Stemcell). Media was changed every 24 hours. Confluency of iPSCs was assessed every 24h for 8 consecutive days using Live/dead staining cell imaging kit (R37601, ThermoFisher) according to manufacturer’s protocol. Live imaging of whole wells was performed with Opera Phenix High Content Imaging System (PerkinElmer) at 20× with three wells per condition and time point. Live cell area was measured by absolute threshold and expressed as percentage of total well area (confluency). Non-linear regression (logistic growth) was performed (GraphPad Prism 10.2.2.).

### Flow cytometry

iCE-BECs with ApoE3 or ApoE4 genetic variant (80.000/well in 24-well plate) were treated with fluorescently labeled transferrin (T13342, ThermoFisher) at 25 µg/mL for 20 min at 37°C or left untreated, followed by incubation of CD31-AF700 (NB600-562AF700 Novus Biological, 1:100) for 30 min at 4°C. Alternatively, iEC and iCE-BECs were collected, fixed in 4% PFA for 30min and incubated with CD31-AF700 (NB600-562AF700, Novus Biological, 1:100) and Claudin-5 (352588, Thermo Fisher, 1:100) for 30 min at 4°C. Cells were washed with FACS buffer (PBS + 0.5% BSA + 2 mM EDTA), and fluorescence was immediately acquired using a Flow Cytometer (CytoFLEX LX, Beckman Coulter) in duplicates. Cells were gated on FSC and SSC. Unstained negative controls were used to adjust gain for FITC or AF700, respectively. Per sample, 10.000 live cells (singlets) were acquired and mean fluorescence of transferrin, PECAM-1 or Claudin-5 was calculated.

### Live imaging of FerroOrange to assess labile iron pool

Intracellular labile iron was measured using BioTracker^TM^ FerroOrange Live Cell Dye, a fluorescent probe that specifically detects labile iron (II) ions (Fe^2+^) only. Briefly, cells were washed and incubated with 1 µM BioTracker^TM^ FerroOrange Live Cell Dye (SCT210, Sigma) and cell marker (CytoTrace Green, 22017, AAT Bioquest) in HBSS at 37°C for 30min. Live cell imaging was performed using Opera Phenix High Content Imaging System (PerkinElmer) at 63×, 4 wells per condition, 10 fields per well, 8 z-stacks à 1 µm. Sum intensity of FerroOrange was normalized to the cell area (absolute threshold of CellTrace). To artificially increase LIP, cells were treated with an iron donor ferrous ammonium sulfate at 100 µM (FAS, 203505, Sigma), while treatment with iron chelator PIH at 10 µM (ab145871) was used to deplete LIP before FerroOrange Live cell dye was applied.

### Calcein-AM assay to assess labile iron pool

Intracellular labile iron was measured using the metal-sensitive probe Calcein acetoxymethyl ester (Calcein-AM), as previously described ^77^. This is a non-fluorescent dye that becomes fluorescent after enzymatic modification once it permeates the cell membrane ^53,77,78^. This fluorophore binds iron stoichiometrically, which quenches its green fluorescence. In short, cells were washed, incubated with 0.5 μM of Calcein-AM (C1430, ThermoFisher) in HBSS and whole cell marker (HCS CellMask™ Stain Deep Red, H32721, ThermoFisher) for 20 min at 37°C. Iron chelator PIH at 10 µM (ab145871, abcam) was applied to half of the wells for 10 min. Cellular calcein fluorescence was measured in live cells using Opera Phenix High Content Imaging System (PerkinElmer) at 20×, 5 wells per condition, 16 fields per well, 3 z-stacks à 1 µm. The ratio between the sum intensity of calcein within the cell area (absolute threshold of CellMask) in untreated cells and iron chelator-treated cells was calculated, reflecting the amount of the labile iron pool. Fold changes were calculated by normalization to ApoE3 per each experiment.

### Statistical analysis

Statistical analyses were performed using GraphPad Prism 10.2.2 software. Normality of the data was evaluated by the Shapiro-Wilk test. For normally distributed numeric data, two-way ANOVA with Sidak’s multiple comparison test was used to compare multiple variables between groups or Student’s t test for comparisons between two groups. For others, non-parametric Mann Whitney U test was used for comparisons between two groups. Detailed statistics are indicated in each figure legend. The data are reported as mean ± SD.

## Acknowledgements

We thank Dr. Sybille Seiler, Dr. Urs Langen and Dr. Colette Bichsel for scientific discussions and suggestions. We would like to thank Giacomo Valsecchi, Lena Jutz, Dr. Shane Clerkin, Antoine Rizkallah, Petra Stäuble, Telma Lopes, Lorena Fabella, Dr. Heloise Ragelle, Alena Spielmann and Pamela Strassburger for excellent technical support. We would like to thank Dr. Udo Hetzel and Barbara Prähauser for their support with the transmission electron microscopy.

## Author contributions

L.B. contributed to project administration, conceptualization, data curation, methodology, investigation, visualization, writing—original draft, and writing—review & editing. N.S.-R. contributed to conceptualization, supervision, writing—review & editing. C.S. contributed to conceptualization, supervision, data curation, methodology, investigation, writing—review & editing. A.A., B.H., L.D.A., K.S., D.V.T., J.F-P., Chi.Z., Chr..Z., S.G. contributed to data curation, methodology, investigation and writing—review & editing. L.F. contributed to conceptualization, writing—review & editing, A.O. contributed to writing— review & editing, M.P. contributed to project administration, supervision, conceptualization, data curation, methodology, investigation, visualization, writing—original draft, and writing—review & editing and decision to submit, R.V. contributed to project administration, supervision, conceptualization, data curation, methodology, visualization, writing—original draft, and writing—review & editing and decision to submit.

## Abbreviations

AD: Alzheimer’s disease
ApoE: Apolipoprotein E
BBB: blood-brain barrier
Calcein-AM: calcein-cetoxymethyl ester
Ct: cycle threshold
DMT-1: Divalent metal transporter 1
FAS: Ferrous ammonium sulfate, an iron chelator
FTH: Ferritin heavy chain
FTL: Ferritin light chain
iCE-BEC: Inducible pluripotent stem cells via Chemical cocktail and Extracellular matrix support-Brain Endothelial Cells
iEC: inducible Endothelial cells
iEC-rep: inducible Endothelial cells treated with Repsox
iPSC: induced pluripotent stem cell
LIP: labile iron pool
NT IgG: non-targeting IgG
P_app_: apparent permeability
PBS: Phosphate buffered saline
PC: principal component
PCA: principal component analysis
PIH: pyridoxal isonicotinoyl hydrazone
RT: room temperature
TBS-T: Tris-buffered saline with 0.1% Tween® 20 Detergent
TfR: Transferrin receptor
TJ: Tight junction

