## Supplemental material for "ApoE4 disrupts intracellular trafficking and iron homeostasis in an improved iPSC-based model of human brain endothelial cells"

**Supplementary Material**

**Supplementary Table 1. Related to Figure 1D.** List of genes for benchmarking iCE-BECs towards endothelial transcriptomic signature. List adapted from Lu, Houghton, Magdeldin, Durán, Minotti, Snead, Sproul, Nguyen, Xiang, Fine, Rosenwaks, Studer, Rafii, Agalliu, Redmond and Lis [26].

| Positive PC1 Loading | | | | | Negative PC1 Loading | | |
|---|---|---|---|---|---|---|---|
| **PECAM1** | KLHL6 | GGT5 | RGCC | ULBP2 | CLDN6 | WDR86-AS1 | ST6GAL2 |
| **CD93** | FAM107A | C22orf34 | HOXA11 | PAPSS2 | GPC3 | TENM3-AS1 | OXTR |
| **MMP1** | TLR4 | NFIA | TNFSF4 | IFIT2 | LIN28A | NSUN7 | NSG1 |
| **MMRN1** | MEG3 | PREX1 | FAM241A | TNFRSF1B | PTN | MEIS3 | PCAT14 |
| **CLEC14A** | EVA1C | HOXA9 | CNTNAP3B | MTSS1 | DSG2 | MYL7 | LRRTM4 |
| **ROBO4** | SELP | MLKL | LHX6 | VASH1 | IGDCC3 | PARP8 | DCDC2 |
| **SRGN** | TNFAIP8L3 | COL8A1 | AFAP1L1 | SORBS2 | CRABP2 | HAPLN1 | STXBP6 |
| **GIMAP4** | NOVA2 | SEMA6B | MPP4 | A2M | DSC2 | WNT5B | CA3 |
| **GIMAP6** | KDR | SYNPO | MLIP | FAM124A | AP1M2 | OVOL2 | SERPINF1 |
| **DIPK2B** | CARD6 | TFEC | SENCR | KLF9 | EPCAM | CCDC144NL-AS1 | PLD5 |
| **ERG** | MYRIP | COL13A1 | GNGT2 | NRG1 | DMKN | NEO1 | IRS1 |
| **VWF** | CXCL8 | PLCL1 | ZFYVE28 | SSTR1 | RBM47 | PCDH11X | SRGAP3 |
| **LAMA4** | ITGB3 | CASP1 | NFIC | TLE2 | ACTC1 | SORL1 | CHMP4C |
| **CLDN5** | MANCR | HTR2B | EMP1 | MEDAG | QPRT | CPVL | LSR |
| **CAVIN2** | CXCR4 | ERAP2 | PARVB | MIR155HG | SPP1 | MXRA8 | AFAP1L2 |
| **STAB1** | DLL4 | CCRL2 | NTSR1 | SLC17A9 | GPC6 | C4orf19 | ADAMTS2 |
| **GIMAP8** | PDE4B | TOX2 | IFI27 | PLA2G4C | SLC1A3 | LINC02381 | CLDN10 |
| **EMCN** | FAM43A | LINC01116 | HAGLR | GIPC3 | TRIM71 | HAS2 | BOC |
| **CD34** | PDE7B | THBS1 | ANKRD55 | IL7R | LIN28B | PROM1 | LINGO1 |
| **ICAM2** | RASGRP3 | PLSCR4 | SH2B3 | SLC43A1 | MSX2 | LEF1 | SCG3 |
| **BCL6B** | PIK3CG | IL15RA | CD109 | PCDH10 | ENPEP | CADM1 | COL4A6 |
| **BMX** | CLDN11 | CRIP2 | CEACAM21 | APOLD1 | COL1A1 | GPRC5C | COL1A2 |
| **FAM124B** | LNCOG | PDE2A | ZEB2 | SYT11 | SERPING1 | SALL4 | FUT9 |
| **LYVE1** | NRN1 | IRAK3 | NTN4 | ZNF469 | ZFP42 | FAM169A | ADCY10 |
| **ACVRL1** | IL33 | ITGA10 | EVI2B | TLL1 | MMP9 | MOB3B | NPFFR2 |
| **ESAM** | IFI44 | GAB3 | PREX2 | LINC01094 | NKAIN4 | CCDC8 | L1CAM |
| **TIE1** | KLF2 | ADCY4 | LINC00520 | FOXF1 | WFDC2 | UNC5C | PKDCC |
| **APLN** | CARD16 | LINC01358 | TNFRSF11A | ST8SIA4 | LCP1 | SH2D4A | TBX3 |
| **PPP1R16B** | MGP | ACE | NFIA-AS2 | UAP1L1 | ROR2 | GRIP1 | FLRT3 |
| **SHE** | INSYN2B | CD163L1 | ABLIM3 | ENTPD1 | ID4 | RIPOR2 | LINC00648 |

| | | | | | | | |
|---|---|---|---|---|---|---|---|
| **ANPEP** | SGIP1 | FRMD3 | KCNN3 | FAM155A | CRYBG2 | DACT1 | IGDCC4 |
| **ARHGEF15** | FERMT3 | SEMA3G | DYSF | USHBP1 | HPGD | ALPK2 | DLK1 |
| **HHEX** | HCLS1 | LONRF3 | SYNE3 | ARHGAP22 | WDR86 | SEMA5A | CORO2A |
| **TAL1** | TMEM204 | CFAP54 | EFEMP1 | ITGA5 | SOX9 | RAMP1 | EDNRA |
| **PTX3** | BMP6 | MYCT1 | WSCD1 | RFLNB | LYPD6B | GLI2 | PLPPR3 |
| **RHOJ** | SPAAR | THBD | EDN1 | LPAR6 | DSP | NRK | PATJ |
| **LINC01235** | GMFG | PD | PKD1L1 | CUBN | APOE | CAMK1G | SALL1 |
| **SOX18** | LDB2 | CHCHD2 | PALD1 | ARAP3 | DPPA4 | CDX2 | |
| **MFNG** | NOTCH4 | CXCL1 | IL18R1 | CSGALNACT1 | EFS | LUM | |
| **GPR4** | TMEM255B | SH3RF3 | VIM | PGF | H2AFY2 | ALPL | |
| **ADGRF5** | CDH5 | MEF2C | PROCR | NID1 | CD24 | PCDHA12 | |
| **PCAT19** | RASIP1 | SHANK3 | COX7A1 | PARP12 | H19 | JPH2 | |
| **GIMAP7** | CHST1 | FLT1 | IL3RA | PMP22 | FBLN1 | IGFBP3 | |
| **IL1RL1** | SCARF1 | NLRC5 | CLEC1A | ST6GALNAC4 | EMB | SOX11 | |
| **FABP4** | CYTL1 | CASP4 | PCSK1 | LINC01197 | NPPB | PCSK1N | |
| **PALMD** | FGD5 | NOS3 | NAV3 | TEK | GPC4 | PKP2 | |
| **ZEB1** | FOXC2 | EGFL7 | INKA1 | SP100 | MARVELD3 | TMEM92 | |
| **MMRN2** | TMEM273 | SAMD9 | CCL2 | PLXND1 | TINCR | XKR4 | |
| **GIMAP1** | P4HA3 | VAMP5 | HIC1 | IFIT3 | TNC | PPP2R2B | |
| **CNRIP1** | FLI1 | PLXNA4 | RAMP2 | NRIP3 | SYTL1 | LINC01224 | |
| **ESM1** | SH3TC1 | MILR1 | NT5E | HRH1 | KRT8 | CNN1 | |
| **TM4SF18** | LGALS1 | CPT1A | TMEM156 | ECE1 | PDPN | SLC4A4 | |
| **GIMAP2** | TDRD10 | CAV1 | HHIP-AS1 | SH3TC2 | NLGN4X | TEAD3 | |
| **HHIP** | RAC2 | KANK3 | TM6SF1 | DYRK3 | IGFBP5 | TRMT9B | |
| **PLVAP** | WDFY4 | ZNF366 | MIR137HG | SYNM | SHANK2 | DSC3 | |
| **ADGRL4** | FAM78A | LRRC70 | NPR1 | LTBP2 | GPR87 | GALNT17 | |
| **ABI3** | NPAS2 | SLFN11 | HOXD9 | DPYD | MFAP5 | MPPED2 | |
| **TNFRSF14** | GBP4 | SERPIND1 | DDR2 | CDKN2C | ARSI | IGF2-AS | |
| **APOL3** | NRGN | SAMSN1 | STX11 | HOXD1 | TRIM55 | ADGRV1 | |
| **LAPTM5** | ADAMTSL1 | KCTD12 | PTPRE | SPESP1 | LRRN4 | CA2 | |
| **PCDH12** | C2CD4B | ANXA2R | DMTN | STEAP1 | RUBCNL | DIO3OS | |
| **ECSCR** | MAPK11 | VEPH1 | STC1 | CLDN14 | PKIB | TNFRSF19 | |
| **ENG** | HLX | LINC02454 | MCTP1 | ADGRA2 | PRSS16 | GAS7 | |
| **SOX17** | HOXD8 | CTSS | LGALS9 | EBF1 | GYG2 | GALNT3 | |
| **TNFSF10** | GRAP | LOX | GBP1 | ITGA11 | PURPL | ERP27 | |
| **GNG11** | NFIB | TBX18 | LRRC8C | MSRB3 | CHPF | KRT19 | |
| **CALCRL** | CPNE5 | STK32B | MIR217HG | CCNA1 | ACSS3 | EPS8L2 | |
| **TM4SF1** | GRASP | MEOX2 | ARHGAP20 | RFTN2 | ADAMTS19 | NR6A1 | |
| **S1PR1** | ARHGEF28 | SPOCK1 | KIAA1549L | ARHGEF6 | CTSV | NTRK2 | |

| | | | | | | |
|---|---|---|---|---|---|---|
| **LYL1** | MALL | RNASE1 | GBP2 | MAMLD1 | RARRES2 | WWC1 |
| **BGN** | LY96 | HOXA10 | MT2A | TBXA2R | SPINT2 | PLBD1 |
| **TMEM173** | GJA4 | BACE2 | UBA7 | IL4I1 | CDH8 | PDGFRB |
| **CLEC2B** | DKK1 | TCTEX1D1 | TGFBR2 | GAPLINC | TFAP2A | LMOD1 |
| **ANGPT2** | STEAP1B | TNS2 | F2RL2 | HSD17B2 | ERVH48-1 | RGS16 |
| **SH2D3C** | THSD1 | LMO2 | TSPO | RAPGEF5 | SLC27A6 | RIMS2 |
| **DOCK10** | LIX1L | NEGR1 | AHNAK2 | IL6R | IGF2 | SSC4D |
| **LINC01013** | CDYL2 | SERPINE1 | NRG3 | OAF | MLPH | PRKCZ |
| **PTPRB** | FLT4 | SNED1 | SLC9A3R2 | ARHGAP24 | MYO5B | FERMT1 |
| | | | ALDH1A2 | NOX4 | | |

*Extraction of the top 500 genes with strongest positive and 500 genes with strongest negative contribution to PC1 from sheet "Fig.1C PC1 Loading Genes" from dataset S02 of the meta analysis fromLu, Houghton, Magdeldin, Durán, Minotti, Snead, Sproul, Nguyen, Xiang, Fine, Rosenwaks, Studer, Rafii, Agalliu, Redmond and Lis [26]. Subset of extracted genes that were expressed in our cells (392 for positive PC1 loading, 193 for negative PC1 loading) are listed below. Genes with positive PC1 loading are associated with endothelial signature, while negative PC1 loading are related to epithelial identity, see Figure 1D*.

Supplementary Figure S1.

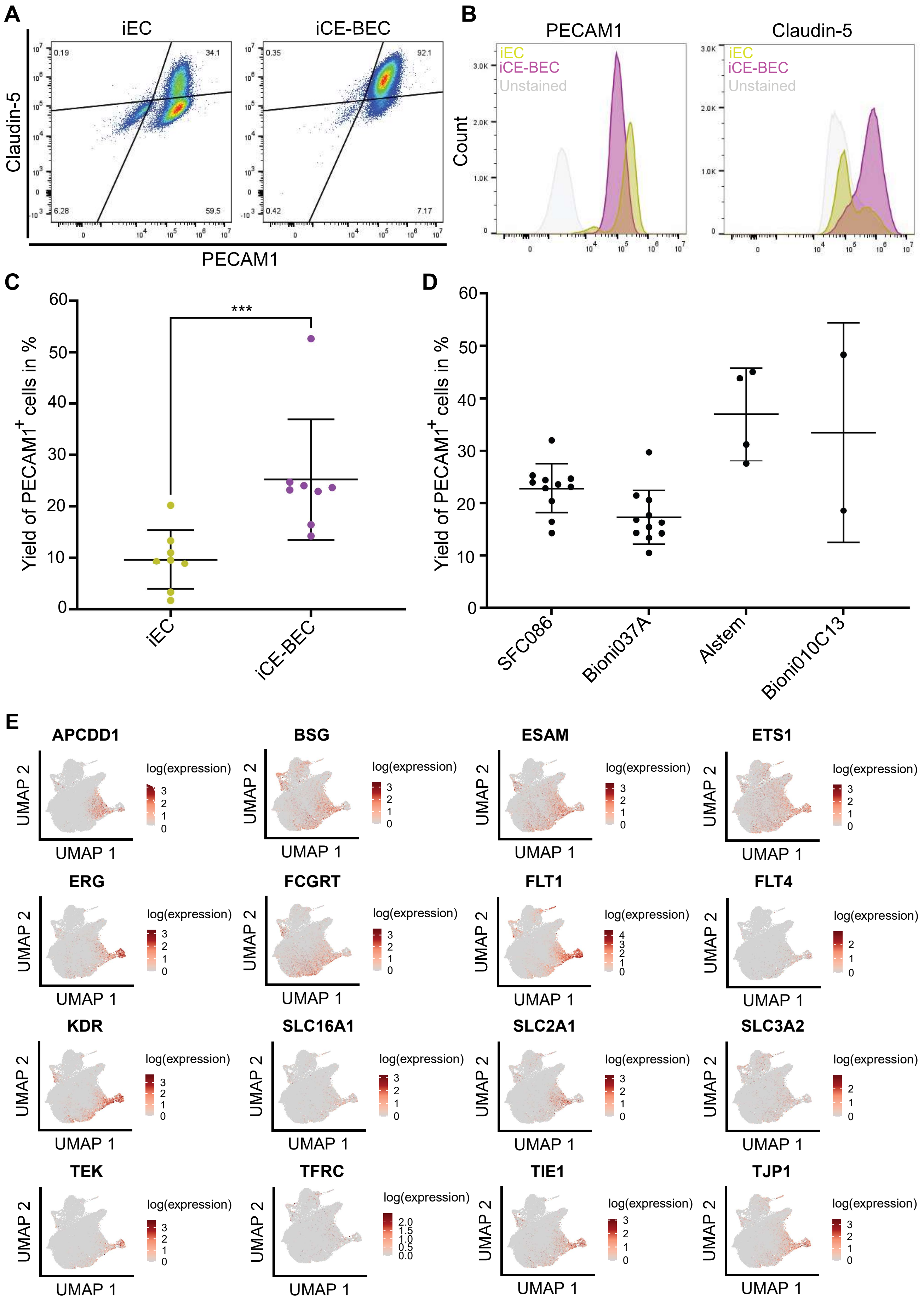

### Supplementary Figures

**Supplementary Figure 1. Related to Figure 1. A,** Dot plot showing flow cytometry assessment of iECs and iCE-BECs on day 14 in culture. The fluorescence intensity PECAM1 and Claudin-5 are plotted on x-axis and y-axis, respectively. Quadrants were set based on unstained controls. Each dot represents an individual cell. The data represent the distribution of PECAM1 and Claudin-5 expression from one independent experiment. **B,** Histograms show fluorescence intensities of PECAM1 (left panel) and Claudin-5 (right panel) on the x-axis and the number of cells on the y-axis from cells plotted in panel A. iECs are shown in yellow, iCE-BECs in magenta and unstained control in gray. **C,** Yield of PECAM1-positive cells before and after MACS sorting at day 11 comparing protocols summarized in Figure 1A using the SFC086 line**.** Each data point represents one differentiation, data come from $n = 7$ individual differentiations. Differences in the yield between iECs and iCE-BECs are statistically significant as evaluated by the Mann-Whitney-U test. ***, $p < 0.001$ **D,** Yield of PECAM1-positive cells before MACS sorting and after MACS sorting at day 11 across different hiPSC lines using the protocol for iCE-BECs. Each data point represents one differentiation. Data come from $n = 11$ individual differentiations for SFC086 and Bioni037A line, $n = 4$ for Alstem line, and $n = 2$ for Bioni010C13 line. **E,** Feature plots showing normalized log expression of marker genes of endothelial and mural markers, plotted on the UMAP from Figure 1C.

Supplementary Figure S2.

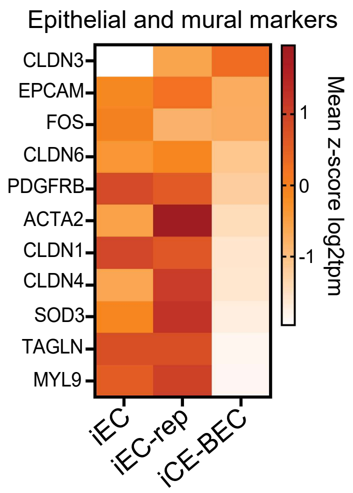

**Supplementary Figure 2. Related to Figure 1.** Bulk RNA-Seq heatmap showing expression of epithelial and mural markers across the three differentiation protocols, iEC, iEC-rep, and iCE-BECs. Values are expressed as mean z-score $log_2$tpm, with three independent differentiations per condition.

Supplementary Figure S3.

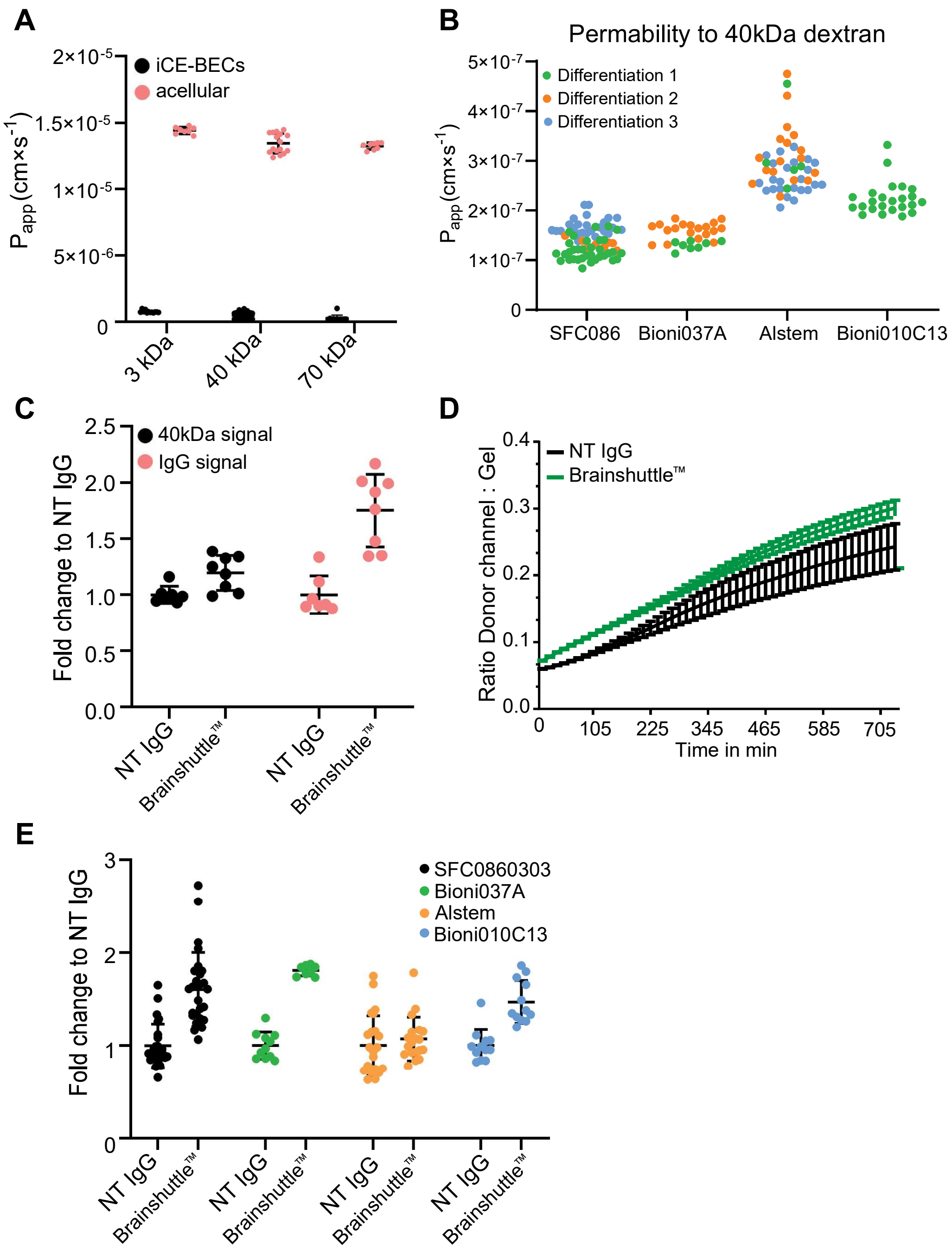

**Supplementary Figure 3. Related to Figure 2**. **A,** Quantification of apparent permeability ($P_{app}$) of iCE-BECs to 3, 40 or 70 kDa dextrans in comparison to the acellular control ("acellular"). Graph shows mean ± SD of one differentiation with 8 technical replicates per condition. **B,** Quantification of apparent permeability ($P_{app}$) of iCE-BECs to 40 kDa dextran using MIMETAS OrganoPlate® 2-lane 96 in iCE-BECs generated with different parental iPSC. Graph shows data from multiple differentiations, each data point represents one chamber. In particular, SFC086, n = 3; Bioni037A, n = 1; Alstem, n = 2; and Bioni010C13, n = 1. **C,** Quantification of relative IgG transcytosis (IgG signal) and apparent permeability to 40 kDa dextran (40 kDa signal) across iCE-BECs after incubation with 200 nM non-targeting IgG (NT IgG) or a Brainshuttle™ antibody. Antibody and dextran values are measured in the same channels, and are normalized relative to the NT IgG condition. Graph shows mean ± SD of one differentiation with 8 technical replicates per condition. **D,** Representative antibody transcytosis curves across iCE-BECs after incubation with 200 nM non-targeting IgG (NT IgG) or Brainshuttle™ antibody. Images were acquired immediately after incubation for 12 hours and ratio between donor channel and gel channel signals were plotted against time (see methods for details). Each curve shows mean ± SEM data from 8 chambers. **E,** Quantification of relative antibody transcytosis across iCE-BECs generated with different parental iPSC lines after incubation with 200 nM non-targeting IgG (NT IgG) or Brainshuttle™ antibody. Graph shows mean ± SD of multiple differentiations with minimum 8 technical replicates per condition. Each data point represents one chamber. In particular, SFC086, n = 3; Bioni037A, n = 2; Alstem, n = 3; and Bioni010C13, n = 1.

Supplementary Figure S4.

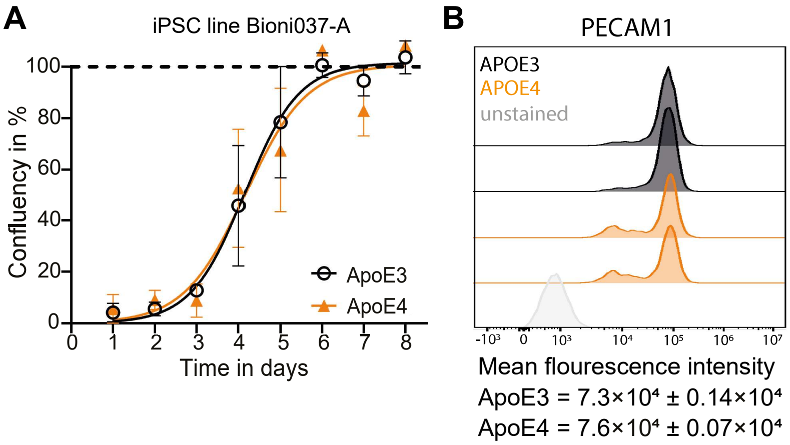

**Supplementary Figure 4 - related to Figure 3. A,** Assessment of proliferation rate of iPSC line Bioni037-A with ApoE3 and ApoE4 genetic variants over eight consecutive days. Live/dead staining and live imaging of whole wells was performed with Opera Phenix High Content Imaging System (PerkinElmer) at 20× magnification with three wells per condition and time point. Live cell area was measured by absolute threshold and expressed as percentage of total well area (confluency) ± SD. Non-linear regression (logistic growth) was performed. **B,** Flow cytometry assessment of PECAM1 in live iCE-BECs with ApoE3 or ApoE4 gene variant. Histograms show fluorescence intensities of PECAM1 on the x-axis and the number of cells on the y-axis. ApoE3 iCE-BECs are shown in black, ApoE4 iCE-BECs in orange and unstained control in gray. Mean fluorescent intensity of PECAM1 in live cells in iCE-BECs with ApoE gene variants from one differentiation, two technical replicates per condition.

Supplementary Figure S5.

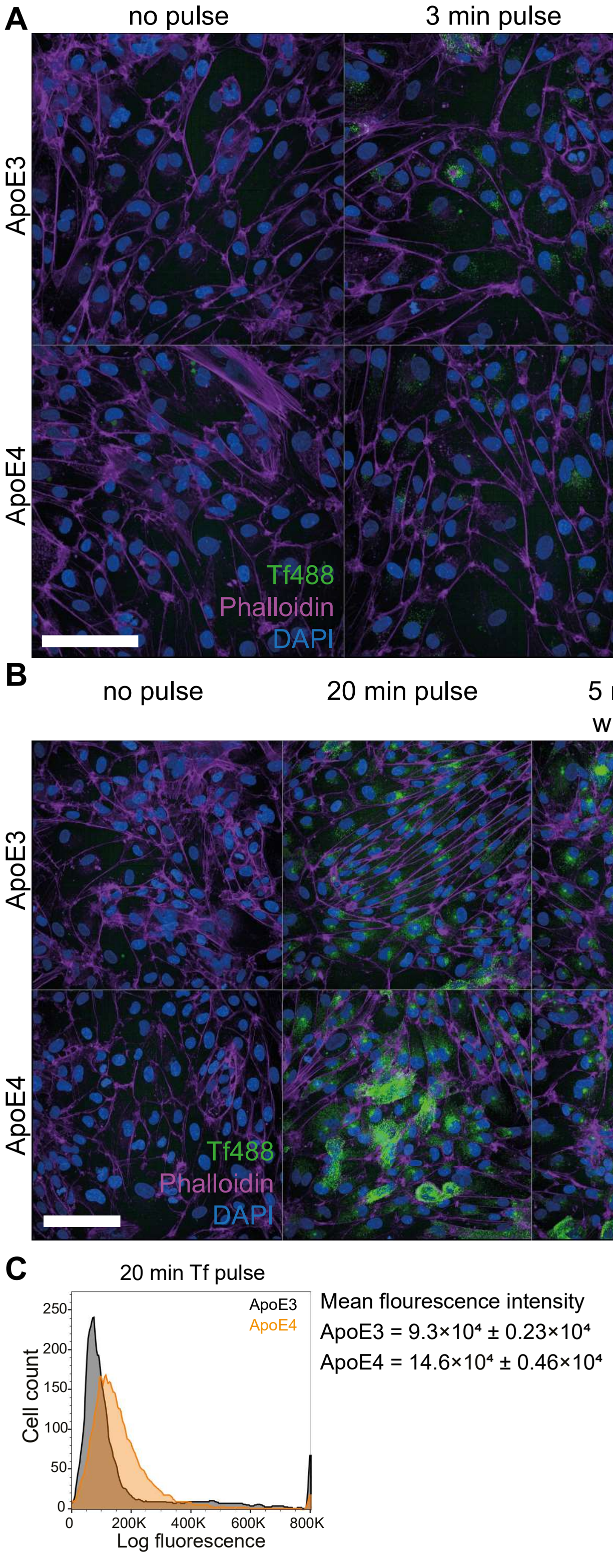

**Supplementary Figure 5 - related to Figure 5. A,** Representative maximum projection confocal images of iCE-BECs with ApoE gene variants from pulse assay and **B,** pulse-chase assay assessing transferrin trafficking kinetics. Briefly, cells were incubated with fluorescently labeled transferrin and fixed after different time points for continuous uptake (pulse assay) while for recycling assessment (pulse-chase assays), cells were incubated for 20 min with fluorescently labeled transferrin followed by incubation of 10-fold higher concentration of unlabeled holo-Transferrin for different time points. Cells are pseudo-colored showing Transferrin in green, Phalloidin in magenta, and DAPI-stained nuclei in blue. Scale bar, 100 µm. For each time point, 50 images were acquired at 40× using a high content screening system, maximal projections were used to quantify sum intensity of transferrin in Phalloidin-positive area. **C,** Flow cytometry assessment of transferrin uptake after 20 min incubation in live iCE-BECs with ApoE3 and ApoE4 genetic variants. Histograms show log fluorescence intensities of fluorescently labeled transferrin on the x-axis and the number of cells on the y-axis. ApoE3 iCE-BECs are shown in black, ApoE4 iCE-BECs in orange. Mean fluorescence intensity ± SD for one differentiation with two technical replicates per condition.

Supplementary Figure S6.

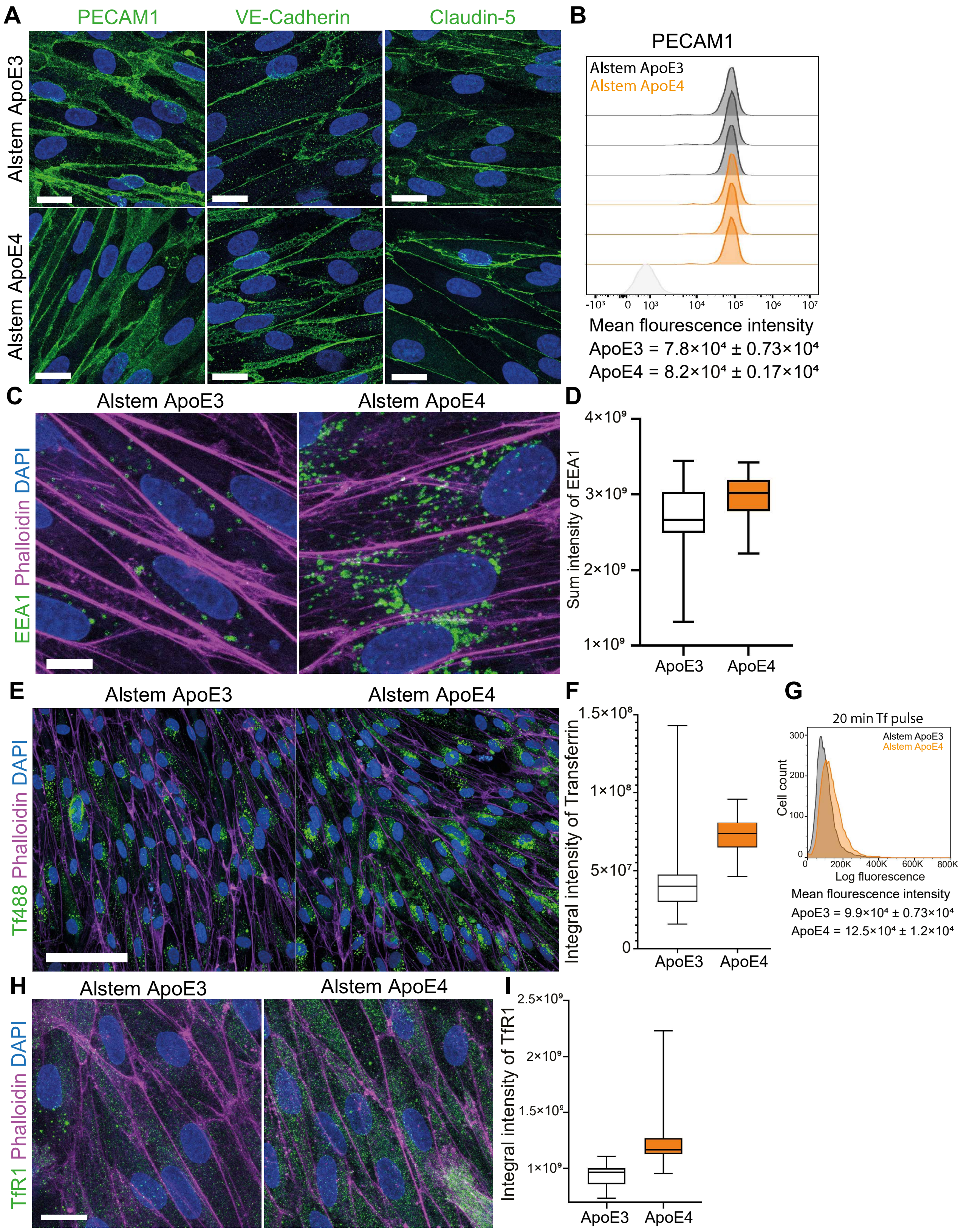

**Supplementary Figure 6 - related to Figure 3, 4 and 5. A,** Representative fluorescence images after immunostaining with endothelial-specific markers iCE-BECs with ApoE3 or APOE4 genetic variant using the Alstem line. Cells are pseudo-colored showing PECAM1, VE-Cadherin or Claudin-5 in green and DAPI-stained nuclei in blue. Scale bar, 20 µm. **B,** Flow cytometry assessment of PECAM1 in live iCE-BECs with ApoE3 or ApoE4 gene variant using the Alstem line. Histograms show fluorescence intensities of PECAM1 on the x-axis and the number of cells on the y-axis. ApoE3 iCE-BECs are shown in black, ApoE4 iCE-BECs in orange and unstained control in gray. Mean fluorescent intensity ± SD of PECAM1 in live cells in iCE-BECs with ApoE gene variants from one differentiation, three technical replicates per condition. **C,** Representative maximum intensity projections of confocal fluorescence images of iCE-BECs with ApoE3 or ApoE4 genetic variant after immunostaining with a marker of early endosomes (EEA1). Cells are pseudo-colored showing EEA1 in green, Phalloidin in magenta and DAPI-stained nuclei in blue. Scale bar, 10 µm. **D,** Quantification of EEA1 sum intensity in Phalloidin-positive area. Graph shows boxplots with interquartile ranges and median. Lines show the 5th and 95th percentiles. Data come from one differentiation, 35 images per condition. **E,** Representative images of iCE-BECs Alstem line with ApoE3 or ApoE4 gene variant pulsed with fluorescently labeled transferrin for 20 min. Cells are pseudo-colored showing transferrin in green, Phalloidin in magenta and DAPI-stained nuclei in blue. Scale bar, 100 µm. **F,** Quantification of integral intensity of transferrin incubated for 20 min normalized to Phalloidin-positive area. Per condition, 50 images have been acquired at 40×, using a high content screening system, maximal projections were used to quantify integral intensity of transferrin in Phalloidin-positive area. **G,** Flow cytometry assessment of transferrin uptake after 20 min incubation in live iCE-BECs with ApoE3 or ApoE4 gene variant. Histograms show fluorescence intensities of fluorescently labeled transferrin on the x-axis and the number of cells on the y-axis. ApoE3 iCE-BECs are shown in black, ApoE4 iCE-BECs in orange. Mean fluorescence intensity ± SD from one differentiation with three technical replicates per condition. **H,** Representative maximum intensity projections of confocal fluorescence images of Alstem iCE-BECs with ApoE3 or

ApoE4 genetic variant after immunostaining with transferrin receptor (TfR). Cells are pseudo-colored showing TfR in green, Phalloidin in magenta and DAPI-stained nuclei in blue. Scale bar, 20 µm**. I,** Quantification of TfR integral intensity in Phalloidin-positive area. Graph shows boxplots with interquartile ranges and median. Lines show the 5th and 95th percentiles, data from one differentiation with 35 images per condition.

Supplementary Figure 7.

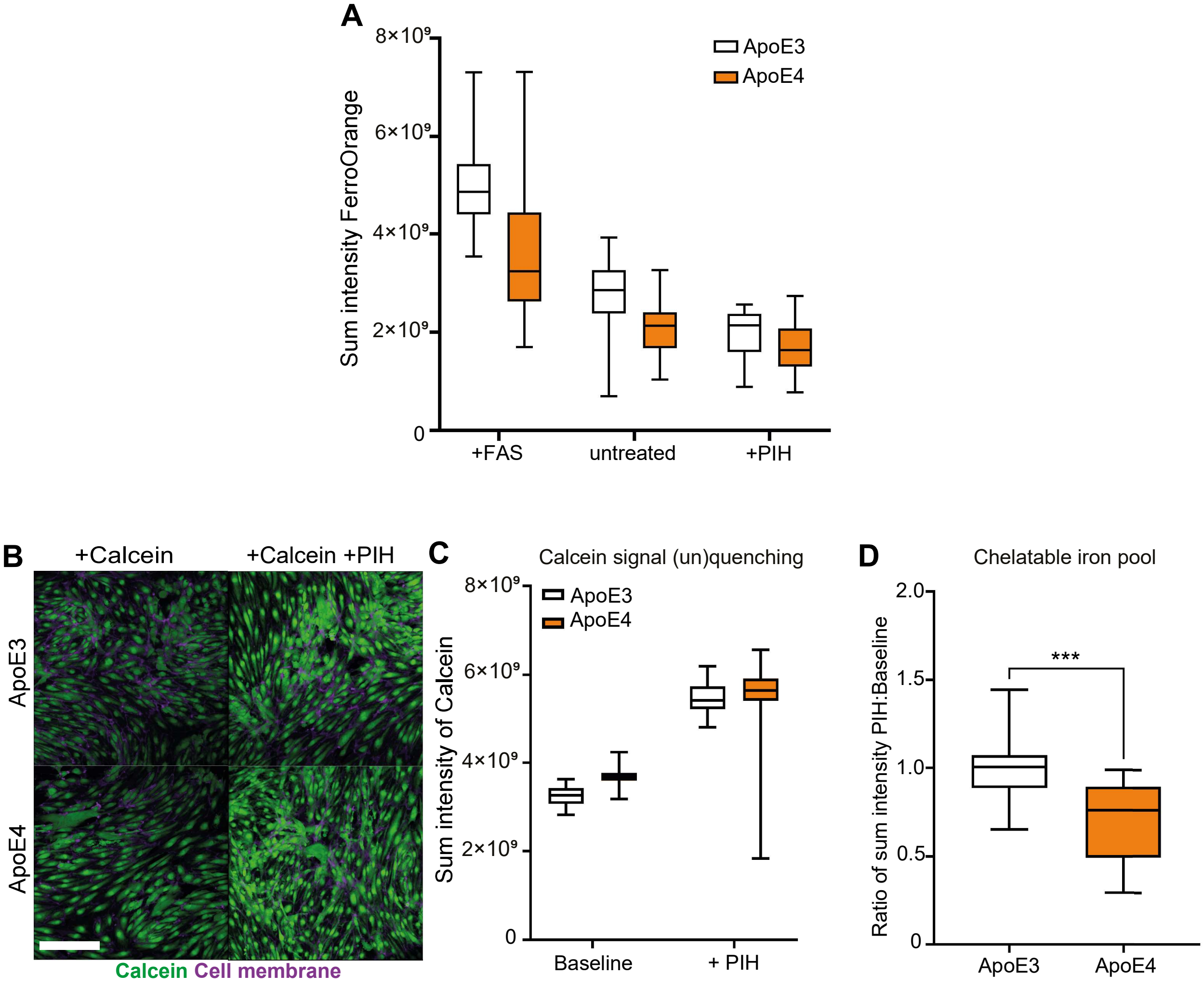

**Supplementary Figure 7. Related to Figure 6. A,** Mean intensity of FerroOrange in iCE-BECs with ApoE3 or ApoE4 gene variant. As controls, cells have been treated with an iron donor ferrous ammonium sulfate (FAS) or iron chelator pyridoxal isonicotinoyl hydrazone (PIH) for 30 min before incubating with FerroOrange, a fluorescent probe that specifically detects labile iron (II) ions ($Fe^{2+}$) in live cells. Graph shows boxplots with interquartile ranges and median. Lines show the 5th and 95th percentiles, data from one differentiation, 40 images per condition have been acquired at 63× using a high content screening system, maximal projections were used to quantify sum intensity of FerroOrange within cells. **B,** Representative images of iCE-BECs with ApoE gene variants incubated with the metal-sensitive probe calcein acetoxymethyl ester (calcein-AM), which quenches its green fluorescence when binding to iron and unquenches upon iron chelator treatment with iron chelator pyridoxal isonicotinoyl hydrazone (PIH). Cells were treated with an iron chelator PIH or left untreated. Cellular calcein fluorescence was measured in live cells using high content screening system at 20×. Cells are pseudo-colored showing Calcein in green, plasma membrane in magenta. Scale bar, 200 µm.**C,** Sum intensity of calcein was normalized per cell area shown in a representative experiment illustrating the calcein signal (un)quenching upon iron chelator (+PIH) treatment. **D,** The ratio between the mean intensity of Calcein within the cell area in untreated cells (baseline) and iron chelator-treated cells (+PIH) was calculated, reflecting the amount of the labile iron pool. Graph shows boxplots with interquartile ranges and median. Lines show the 5th and 95th percentiles, data from n = 3 independent differentiations with 120 images per experiment. Differences in the FerroOrange intensity between ApoE genetic variants are statistically significant as evaluated by the Mann-Whitney-U test ($p < 0.001$).

**Supplementary Video S1 - related to Figure 4.**

Representative video of transferrin (green) intracellular transport in live iCE-BECs with ApoE3 gene variant. Cells were incubated with fluorescently labeled transferrin for three hours and then videos of one minute were acquired at 100× using a Widefield microscope. Representative image frames of those videos are shown in Figure 4E.

**Supplementary Video S2 - related to Figure 4.**

Representative video of transferrin (green) intracellular transport in live iCE-BECs with ApoE4 gene variant. Cells were incubated with fluorescently labeled transferrin for three hours and then videos of one minute were acquired at 100× using a Widefield microscope. Representative image frames of those videos are shown in Figure 4E.
